# An extended and improved CCFv3 annotation and Nissl atlas of the entire mouse brain

**DOI:** 10.1101/2024.11.06.622212

**Authors:** Sébastien Piluso, Csaba Verasztó, Harry Carey, Émilie Delattre, Thibaud L’Yvonnet, Éloïse Colnot, Armando Romani, Jan G. Bjaalie, Henry Markram, Daniel Keller

## Abstract

Brain atlases are essential for quantifying cellular composition in mouse brain regions. The Allen Institute’s Common Coordinate Framework version 3 (CCFv3) is widely used, delineating over 600 anatomical regions, but it lacks coverage for the most rostral and caudal brain parts, including the main olfactory bulb, cerebellum, and medulla. Additionally, the CCFv3 omits key cerebellar layers, and its corresponding Nissl-stained reference volume is not precisely aligned, limiting its utilisability. To address these issues, we developed an extended atlas, the Blue Brain Project CCFv3 augmented (CCFv3aBBP), which includes a fully annotated mouse brain and an improved Nissl reference aligned in the CCFv3. This enhanced atlas also features the central nervous system annotation (CCFv3cBBP). Using this resource, we aligned 734 Nissl-stained brains to produce an average Nissl template, enabling an updated distribution of neuronal soma positions. These data are available as an open-source resource, broadening applications such as improved alignment precision, cell type mapping, and multimodal data integration.

## INTRODUCTION

Reference atlases play a crucial role in supporting the brain understanding (Ng et al., 2007; Hawrylycz et al., 2014; Bjerke et al., 2018; Tward et al., 2020; Perens et al., 2023) in both healthy and pathological subjects. Over the past years, several digital atlases of the rodent brain have been created. A well-known open access atlas for the mouse was produced by the Allen Institute for Brain Science (AIBS) in 2007 (Lein et al., 2007; Lau et al., 2008). Additionally, other atlases have been developed through initiatives by the International Neuroinformatics Coordinating Facility, such as the Waxholm Space atlases of the mouse brain (Johnson et al., 2010; Papp et al., 2014). Each of these atlases has been updated over time, guided by improvements in optical microscopy and brain image analysis (Ragan et al., 2012; Johnson et al., 2015; Amato et al., 2016; Milligan et al., 2019; Voigt et al., 2019; Perens et al., 2023).

Most atlases are based on a limited number of brains (Kovacevic et al., 2005; Ma et al., 2005; Bertrand et al., 2008, Johnson et al., 2010). The AIBS produced a mouse brain reference atlas called the Common Coordinate Framework (CCF), whose latest version (CCFv3) is based on a serial two photon tomography (STPT) average template, derived from a significant number of individuals: 1,675 mouse brains (Lein et al., 2007; Kuan et al., 2015; Wang et al., 2020). Over the last ten years this atlas has become a standard in the mouse neuroscientific community (see Chon et al., 2019). It has encouraged classification and quantification of cells in the brain within this reference space (Kim et al., 2017; Ecker et al., 2019; Erö et al., 2018; Murakami et al., 2018; Iqbal et al., 2019; Chen et al., 2019; Rodarie et al., 2022; Yao et al., 2023; Zhang et al. 2022; Shi et al., 2022; Leergaard and Bjaalie, 2022; Zhang et al. 2023). The CCFv3 atlas’s use has expanded to a wide range of applications in medical brain imaging: in two dimensional (2D) histological slices (Tappan et al., 2019; Puchades et al., 2019; Yates et al., 2019; Song et al., 2020; Carey et al., 2023; Sadeghi et al., 2023; Piluso et al., 2023; Young et al., 2023), in 3D histology data using clarification and light-sheet fluorescence microscopy (Renier et al., 2016; Liebmann et al., 2015; Perens et al., 2021; Perens et al., 2023), and in other fields (Ortiz et al., 2020; Wang et al., 2023).

The CCF is an invaluable resource for the scientific community, especially for studying and modeling neuronal circuits in the brain. These qualities align with the initiative of the *Blue Brain Project*, which endeavors to reconstruct and simulate the entire mouse brain (Markram, 2006; Markram et al., 2015). However, CCFv3 is limited in that it does not cover the entire central nervous system of the mouse, nor the entire brain. Moreover, though the AIBS delivered a registered Allen Reference Atlas (ARA) Nissl version to the average template, this is not perfectly aligned in the CCFv3. In this study, we overcome these limitations, providing a well-aligned Nissl template and a comprehensive atlas of the mouse brain and spinal cord.

The two most recent AIBS’s CCF versions are the CCFv2 and CCFv3 (Wang et al., 2020; **Figure 1**). Each was delineated using a different template volume, with the CCFv2 using the reconstructed Allen Reference Atlas coronal Nissl-stained volume (ARA Nissl_COR_) and the CCFv3 using the STPT average template. They both depict the anatomy of the brain but in different ways. While in general the CCFv3 is much improved compared to the CCFv2, the CCFv2 retains several advantages. The CCFv2 annotation volume (ANNOTv2) includes more labels in the grey matter compared to the CCFv3 annotation volume (ANNOTv3), although these annotations are not reliable in the sagittal and horizontal planes, as they were drawn only in the coronal plane (**Figure 1C**). This restricts the use of this version of the atlas for image segmentation (Erö et al., 2018; Krepl et al., 2021; Rodarie et al., 2022). The ARA Nissl_COR_ is highly valuable for the automated registration of both Nissl and *in situ* hybridization (ISH) stained histology since ISH tissue often has a comparable appearance to Nissl. However, the most rostral and caudal brain coronal slices were removed from the dataset as the outermost slices often exhibit distortion, damage, or artifacts due to the limitations of the slicing and staining process, which can compromise the accuracy and reliability of the data (Dong, 2008, Xiong et al., 2018).

**FIGURE 1.**
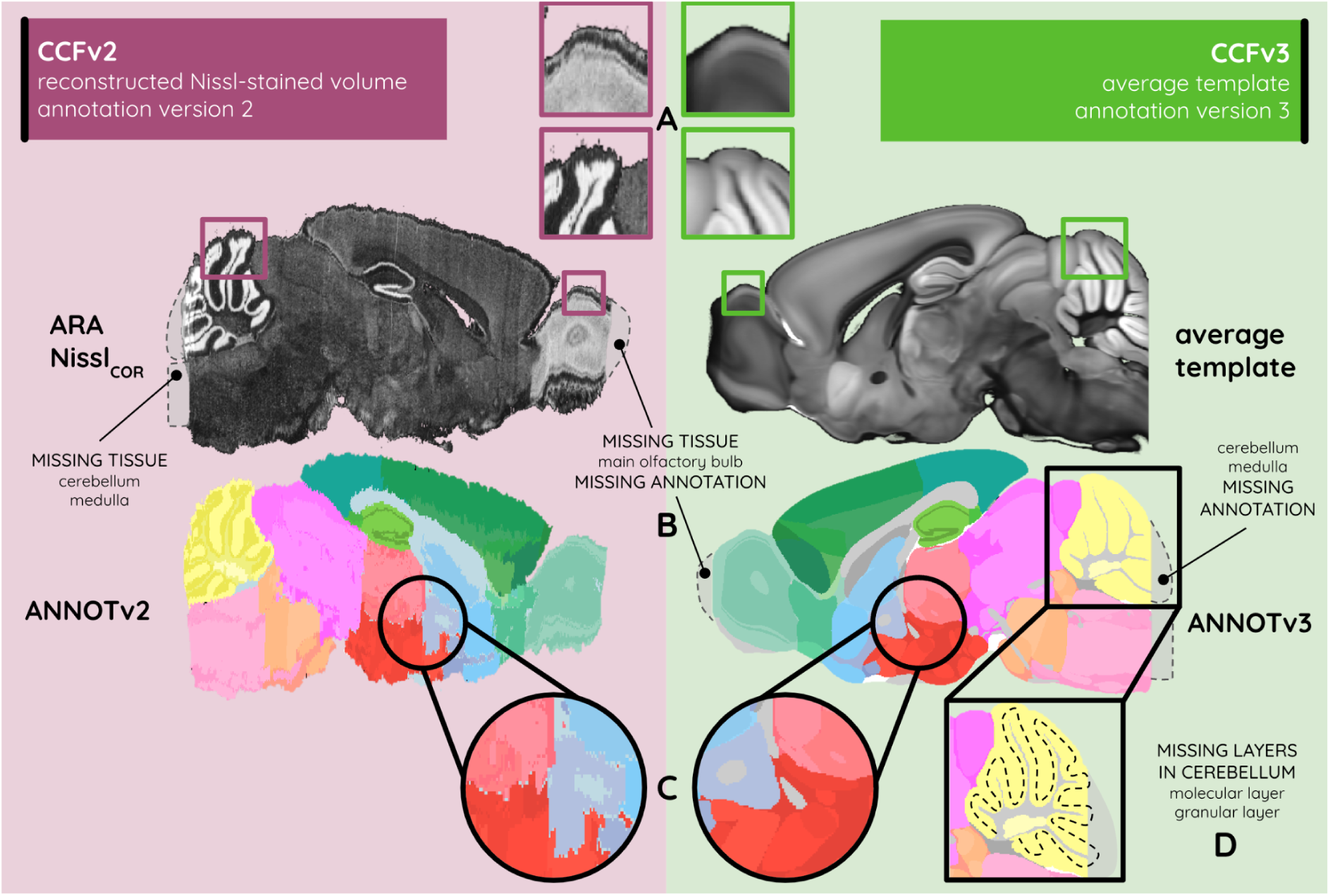
The two latest versions of the atlas provided by the Allen Institute. The CCFv2 (burgundy) and CCFv3 (green) with their respective characteristics: (A) in contrast, texture, and smoothness of the anatomical tissue; (B) in the missing main olfactory bulb, cerebellum, and medulla tissue and annotation; (C) in the 3D smoothness of the annotation; and (D) in subdivisions from the cerebellar annotation.

The result is an incomplete mouse brain for both the ANNOTv2 and the ARA Nissl_COR_, since parts of the olfactory bulb, cerebellum, and medulla are missing (see **Figure 1B**). The cerebellum is especially composed of 16 lobules divided into 3 layers: the granular, the molecular, and the Purkinje layer, the latter being located between the first two. The latest ANNOTv3 version has the molecular and granular layers in the cerebellum missing compared to the ANNOTv2 (see **Figure 1D**). Layers from the latest ANNOTv3 are smooth as they were estimated and validated by experts in 3D using a large multimodal dataset (see **Figure 1C**), enhancing its usability in any orientation as well as for 3D atlas-based segmentation. However, the method used to create the STPT average template is rarely used in laboratories and its choice as a reference has been discussed within the scientific community (Staeger et al., 2020; Rodarie et al., 2022). As a result, scientists must perform multimodal registration between their histological data and the STPT average template if they want these data to be segmented by this version of the atlas. Moreover, while averaging a large number of brains enhances the representativeness of the anatomical reference data, it alters the textures and contrasts in the image partly due to inter-individual variability (see **Figure 1A**). This causes the STPT average template images to diverge even further from the single brain data typically produced by conventional histological platforms.

Single-modality registration, where the template and the data being registered are of the same type, typically yields higher quality results compared to multimodal registration, where the template and data are different types (Goubran et al., 2019; Chiaruttini et al., 2022). The optimal choice of anatomical reference for single-modality registration in many investigations is the ARA Nissl_COR_ (available for CCFv2) while most investigators prefer to use the CCFv3 atlas annotations (ANNOTv3). Thus, a version of the ARA Nissl_COR_ accurately aligned in the CCFv3 space would offer an advantage by allowing single-modality registration combined with the ANNOTv3. While AIBS has attempted to enhance its alignment, the correspondence between the ARA Nissl_COR_ and CCFv3 remains poor, with significant misalignments in the hippocampus, cerebellum, and multiple other regions (see **Figure 5**). In spite of this, the ARA Nissl_COR_ remains frequently used in significant studies (Zhang et al., 2023), potentially leading to inaccurate results as reported by Rodarie et al., 2022. Furthermore, although the CCFv2 initial ARA Nissl_COR_ and the STPT average template were both symmetrical given the inter-hemispheric plane, the resulting pre-registered ARA Nissl_COR_ provided by AIBS in the CCFv3 space is asymmetrical. This further explains the suboptimal precision with which the ARA Nissl_COR_ can be aligned in the CCFv3 space.

The use of 3D digital atlases has grown significantly as scientists seek more precise localization of brain data. An atlas covering the entire brain, without the limitations of histological sectioning, is needed, especially for *in silico* modeling approaches (Markram, 2006; Markram et al., 2015). In particular, growing scientific interest in the olfactory bulb (Kosaka et Kosaka, 2015; Imamura et al., 2020; Licht et al., 2023) and the cerebellum (Golpinar et Demir, 2020; Pisano et al., 2021; Kebschull et al., 2024) means that an atlas covering the entire mouse brain would find immediate application in these regions. Several atlases that incorporate missing regions have been made (Bertrand et al., 2010; Young et al., 2021; Pisano et al., 2021; Wang et al., 2023; Perens et al., 2023) but to our knowledge, no extended CCFv3 version covering the entire central nervous has been constructed.

In this study, we present the CCFv3a_BBP_ Nissl-based atlas which extends the ARA Nissl_COR_ and ANNOTv3 volumes into ARAva_BBP_ Nissl_COR_ and ANNOTv3a_BBP_ to ensure comprehensive coverage of the entire mouse brain based on the CCFv3 (**Figure 2**; see **Video V1** at https://doi.org/10.5281/zenodo.13640418). We identified and aligned complementary Nissl-stained datasets to fill missing regions in the olfactory bulb, cerebellum, and medulla (**Figure 2E-F**). Using a region-focused registration pipeline, we incorporated granular and molecular layers in the cerebellum (**Figure 2G**). Besides, we designed an automated and reproducible method for precisely aligning the ARA Nissl_COR_ into the CCFv3. This approach transformed the multimodal registration challenge into a monomodal problem, allowing for detailed 3D alignment. We further aligned 734 Nissl-stained mouse brains from the Allen Institute database (Lein et al., 2007; Ng et al., 2007) in the CCFv3a_BBP_ to generate an average Nissl template, refined cell distribution placements (**Figure 2E**), and merged recent cortical and spinal cord annotations, resulting in the complete CCFv3c_BBP_ annotation of the mouse central nervous system (**Figure 2D**).

**FIGURE 2.**
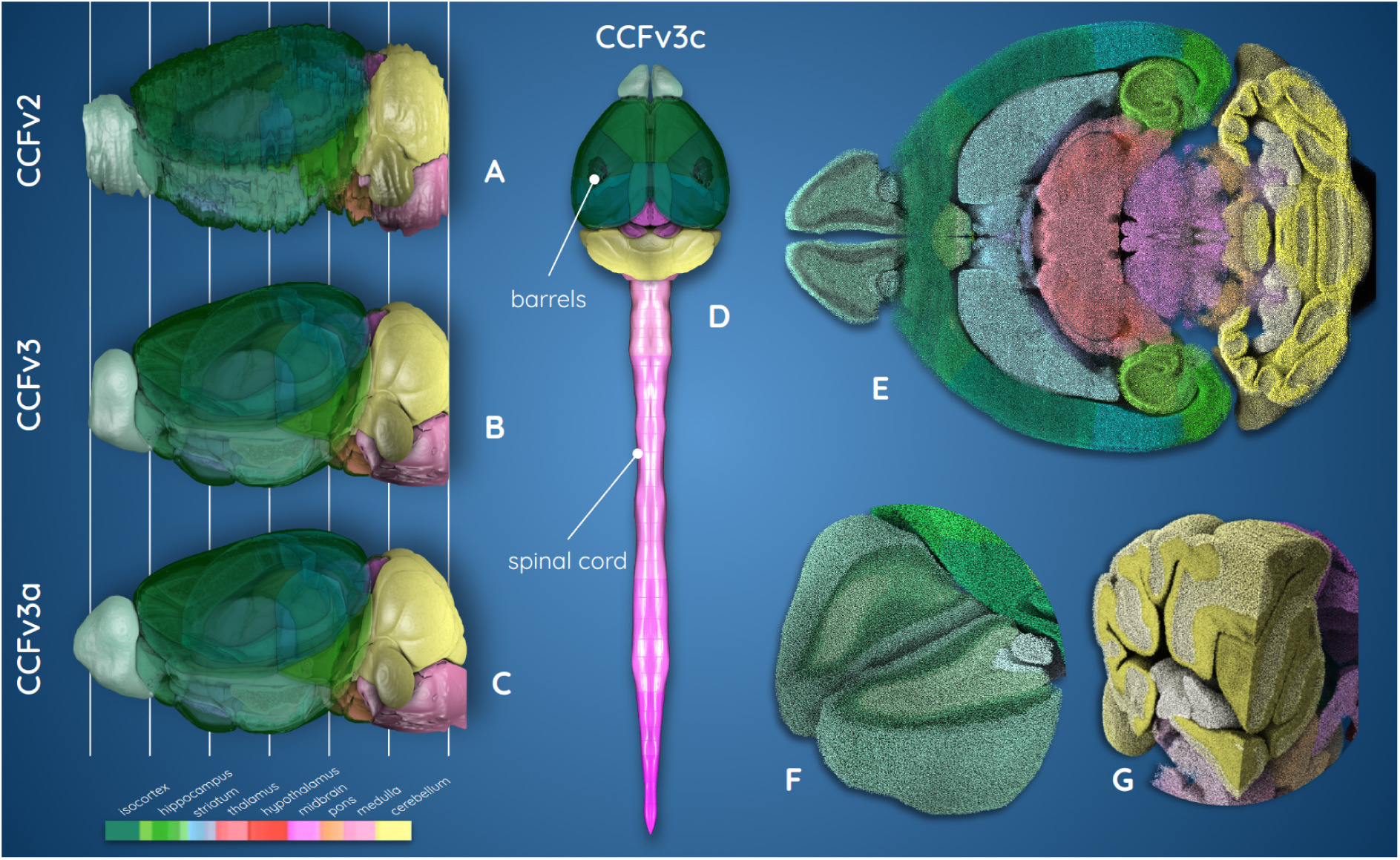
Atlas versions from the AIBS and our extended one (colored according to AIBS annotation color schema). (A) CCFv2 annotation, (B) CCFv3 annotation, (C) our CCFv3a_BBP_ extended annotation, (D) our CCFv3c_BBP_ entire mouse brain version including barrels and the spinal cord, and (E-G) the new cell distribution placement, with each neuron’s soma represented by a sphere, glial cells are excluded here for visualization purposes (E) sliced horizontally, (F) cross section of the main olfactory bulb, (G) cerebellum and medulla (see **Video V1** at https://doi.org/10.5281/zenodo.13640418).

We expect our new atlas to have broad impact on the Neuroscientific community. Other groups have constructed atlases based on the CCFv3 using different modalities such as lightsheet microscopy or MRI (Perens et al., 2023, Wang et al., 2023). Although the templates for these other atlases include the entire olfactory bulb, cerebellum, and brainstem, these regions remain unannotated due to their incomplete representation in the CCFv3. Beyond tools developers, researchers interested in these regions are often forced to either see the closest existing section of the atlas (Zhu et al.,2023), or exclude data which falls outside the atlas boundaries entirely (Yuan et al., 2023). By providing a more accurately aligned Nissl template we have enabled more accurate alignment of data into the CCFv3 and CCFv3a_BBP_. Finally, the existence of whole-brain atlases has enabled multiple large-scale brain-wide analysis projects (Leergaard and Bjaalie, 2022). Here, by providing a continuous atlas of the whole mouse brain and spinal cord, we hope to usher in an era of whole central nervous system analysis.

## MATERIAL AND METHODS

### Source Data

The source data used in this study come from the Allen Institute for Brain Science (AIBS) and can directly be downloaded from its platform (see **Table 1**). The two latest versions are: (1) CCFv2, and (2) CCFv3 (Wang et al., 2020; Allen Institute for Brain Science, 2015). Both versions are available in isotropic resolutions of 25 µm and 10 µm. The original structural annotations are organized hierarchically, and the hierarchy file is the same for both versions. Both annotation datasets, ANNOTv2 and ANNOTv3, have their own number of anatomical regions, which is less than the total number of regions present in the hierarchy. This number varies for each version depending on the experimental data used to produce the annotation file.

**Table 1.** Acronyms, definition, and origin of the data used and produced in this study (see Supplementary Material S1 for a more detailed definition of acronyms, in order of appearance in the text)

| **AIBS** | **Allen Institute for Brain Science** |
| --- | --- |
| ARA Nissl<sub>COR</sub> | Allen Reference Atlas coronal Nissl-stained volume pre-aligned in the CCFv3 by the AIBS |
| ANNOtv2 | Annotations incorporating region delineations defined on the ARA Nissl<sub>COR</sub> |
| STPT average template | Serial Two-Photon Tomography average template |
| ANNOtv3 | Annotations incorporating region delineations defined on the STPT average template |
| CCFv2 = ARA Nissl<sub>COR</sub> + ANNOtv2 | Common Coordinate Framework version 2 |
| CCFv3 = STPT average template + ANNOtv3 | Common Coordinate Framework version 3 |
| AIBS Nissl<sub>SAG</sub> | AIBS sagittal Nissl-stained volume |
| WAXH Nissl<sub>HOR</sub> | Waxholm horizontal Nissl-stained volume |
| **BBP** | **Blue Brain Project** |
| ARA<sub>BBP</sub> Nissl<sub>COR</sub> | ARA Nissl<sub>COR</sub> accurately aligned in the CCFv3 by the BBP |
| ARAv3a<sub>BBP</sub> Nissl<sub>COR</sub> | ARA Nissl<sub>COR</sub> accurately aligned and extended in the CCFv3 by the BBP |
| ANNOtv3a<sub>m</sub> | Expert manual delineation produced by the BBP of the extended tissue based on ARAv3a<sub>BBP</sub> Nissl<sub>COR</sub> as well as of the granular and molecular layers in the cerebellum |
| ANNOtv3a<sub>BBP</sub> | Extended CCFv3 annotation produced by the BBP covering the entire mouse brain plus including new cerebellar layers, assessed using ANNOtv3a<sub>m</sub> |
| CCFv3a<sub>BBP</sub> = ARA<sub>BBP</sub> Nissl<sub>COR</sub> + ANNOtv3a<sub>BBP</sub> | Entire mouse brain atlas derived from CCFv3 produced by the BBP |
| ANNOtv3c<sub>BBP</sub> | Extended CCFv3a<sub>BBP</sub> annotation produced by the BBP covering the mouse central nervous system including spinal cord as well as barrel columns |
| CCFv3c<sub>BBP</sub> = ARA<sub>BBP</sub> Nissl<sub>COR</sub> + ANNOtv3c<sub>BBP</sub> | Mouse central nervous system atlas derived from CCFv3 produced by the BBP including spinal cord and barrel columns |
| average Nissl template | An averaged Nissl-stained template in the CCFv3a<sub>BBP</sub> |
| AIBS<sub>BBP</sub> Nissl<sub>SAG</sub> | AIBS sagittal Nissl-stained volume aligned in the CCFv3a<sub>BBP</sub> by the BBP |
| WAXH<sub>BBP</sub> Nissl<sub>HOR</sub> | Waxholm horizontal Nissl-stained volume aligned in the CCFv3a<sub>BBP</sub> by the BBP |

To reconstruct the missing parts of the ARA Nissl_COR_ reference, we used a sagittally sectioned whole mouse brain from the AIBS (AIBS Nissl_SAG_, Allen Mouse Brain Atlas ID 100042147; Kuan et al., 2015; Allen Institute for Brain Science, 2011). It was downloaded at both isotropic resolutions (see **Supplementary Material S2** for more details). We also made use of another whole mouse brain from the Waxholm space sectioned in the horizontal incidence (WAXH Nissl_HOR_) that we reconstructed in 3D (see **Supplementary Material S2** for more details) with a raw resolution of 9.9 ☓ 21.0 ☓ 9.9 µm^3^ (Johnson et al., 2010). All data correspond to adult C57BL/6J mouse brains.

### Methods

To create an improved annotation and Nissl-based atlas, we firstly ensured data compatibility with some preprocessing steps before applying the registration pipeline we developed (see section 1). This registration pipeline (**Figure 3**) is composed of three automated registration methods (ARM): ARM_A_ for the alignment of the ARA Nissl_COR_ in the CCFv3 (**Figure 4**), ARM_B_ dedicated to the extension of the main olfactory bulb (rostral part of the brain; **Figure 5**), and ARM_C_ dedicated to the extension of the cerebellum and medulla (caudal part of the brain) plus adding the missing cerebellar granular and molecular layers (**Figure 6**). All the Nissl tissue corresponding to the extension added from the AIBS Nissl_SAG_ and the WAXH Nissl_HOR_ was merged into the ARA_BBP_ Nissl_COR_, creating the ARAva_BBP_ Nissl_COR_, in accordance with the AIBS initiative, maintaining a consistent common reference framework. The AIBS Nissl_SAG_ and the WAXH Nissl_HOR_ volumes were independently aligned to the final ARAva_BBP_ Nissl_COR_, as AIBS_BBP_ Nissl_SAG_ and WAXH_BBP_ Nissl_HOR_, to integrate them as additional data into the ANNOTv3a_BBP_. The main ARM_A_ method focused alignment with precision on each region, while simultaneously taking into account the characteristics of all regions. We used the reciprocal combination of the CCF anatomical and annotation files to perform the region selection. Another challenge was the large-scale multimodality, for which we introduced a method that prioritizes region-by-region registration. This approach avoided optimization of a global nonlinear transformation (uses complex transformations to achieve precise alignment by accommodating local deformations) and focused on a set of sub-transformations for each region instead. Based on the ARAva_BBP_ Nissl_COR_ data, experts produced a version of the labels ANNOTv3a_m_ in manually delineating the extended tissue as well as the granular and molecular layers in the cerebellum (see **Supplementary Material S3.3**). Independently, we produced labels for the newly reconstructed tissue in the ANNOTv3a_BBP_ using a semi-automated labeling method including image processing techniques and minimal manual corrections (see **Supplementary Material S3.4**). We used the manual expert annotation ANNOTv3a_m_ for validating the newly added labels in the ANNOTv3a_BBP_ (see **Supplementary Material S3.5**). This segment of the work benefited from the support of the Paxinos and Franklin paper atlas Fifth edition (Paxinos et Franklin, 2019) as an additional atlas reference. We also used the expert delineation of the AIBS Nissl_SAG_ on the web interface from the Allen Institute (Allen Institute for Brain Science, 2011).

**FIGURE 3.**
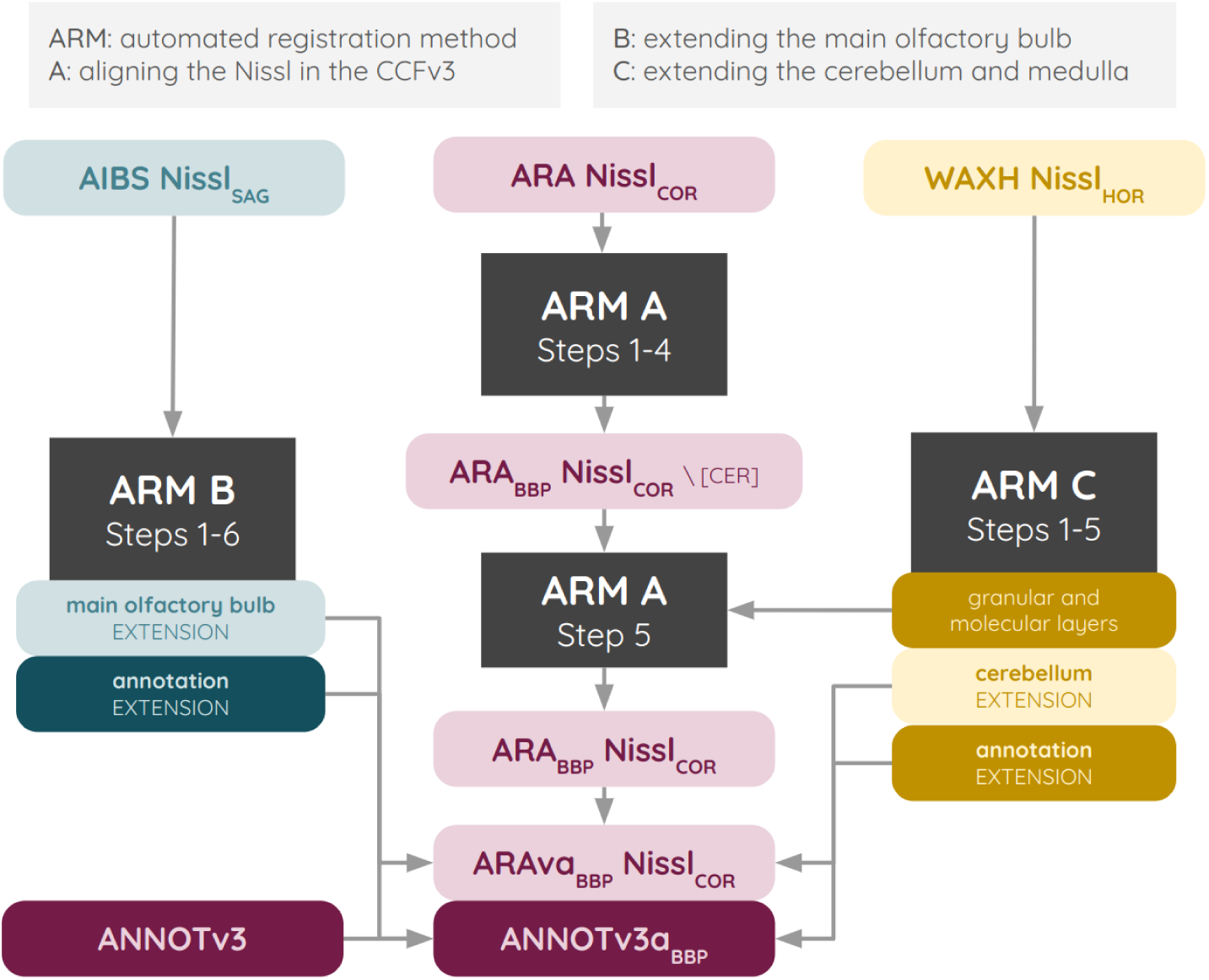
Automated registration pipeline for the precise alignment of the Nissl in the CCFv3 and the extension of the CCFv3, composed of the automated registration methods ARM_A_, ARM_B_, and ARM_C_.

**FIGURE 4.**
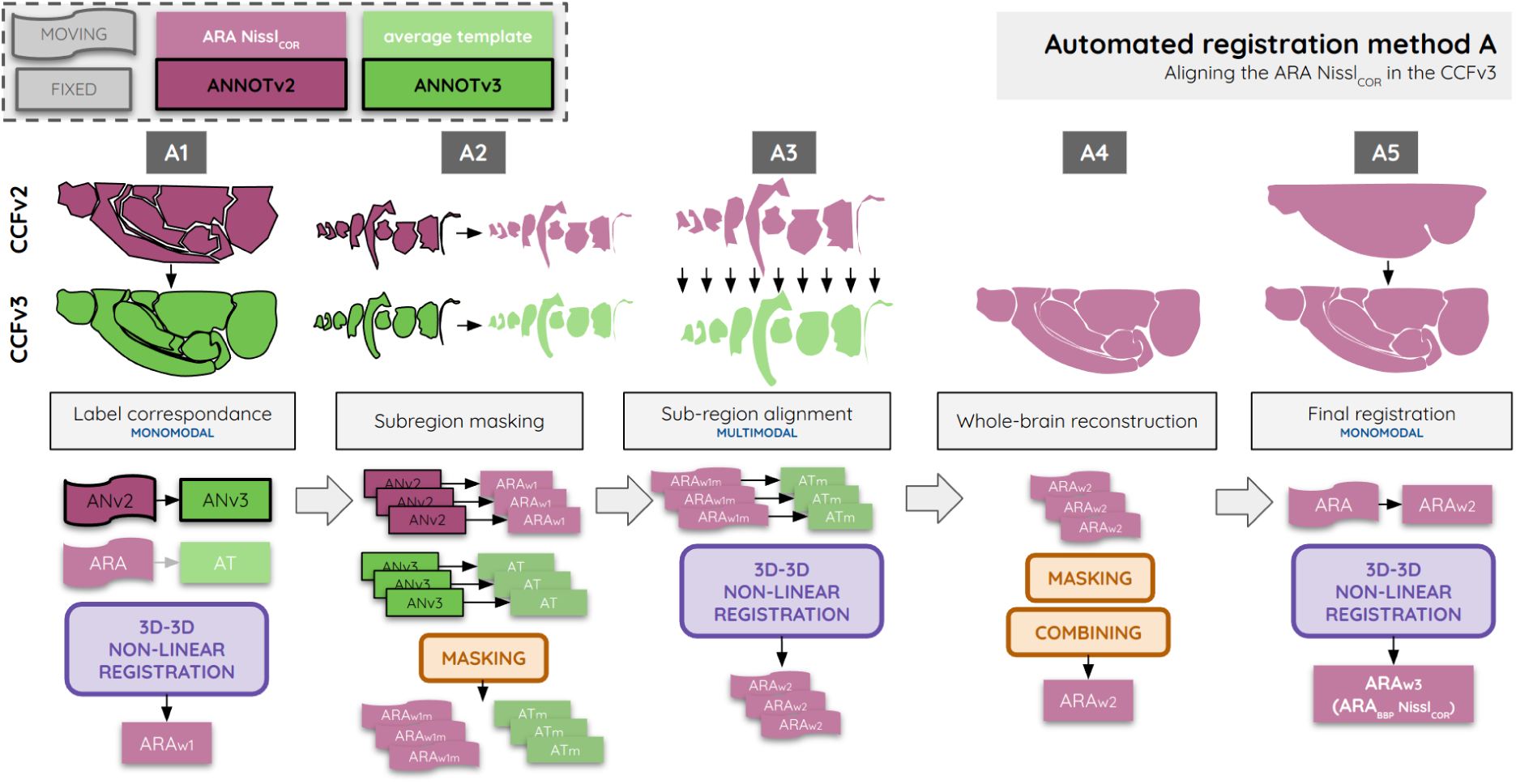
The five-steps region-based guided registration method ARM_A_ for aligning the ARA Nissl_COR_ in the CCFv3. The five steps for the ARM_A_ method are ranging from A1 to A5.

**FIGURE 5.**
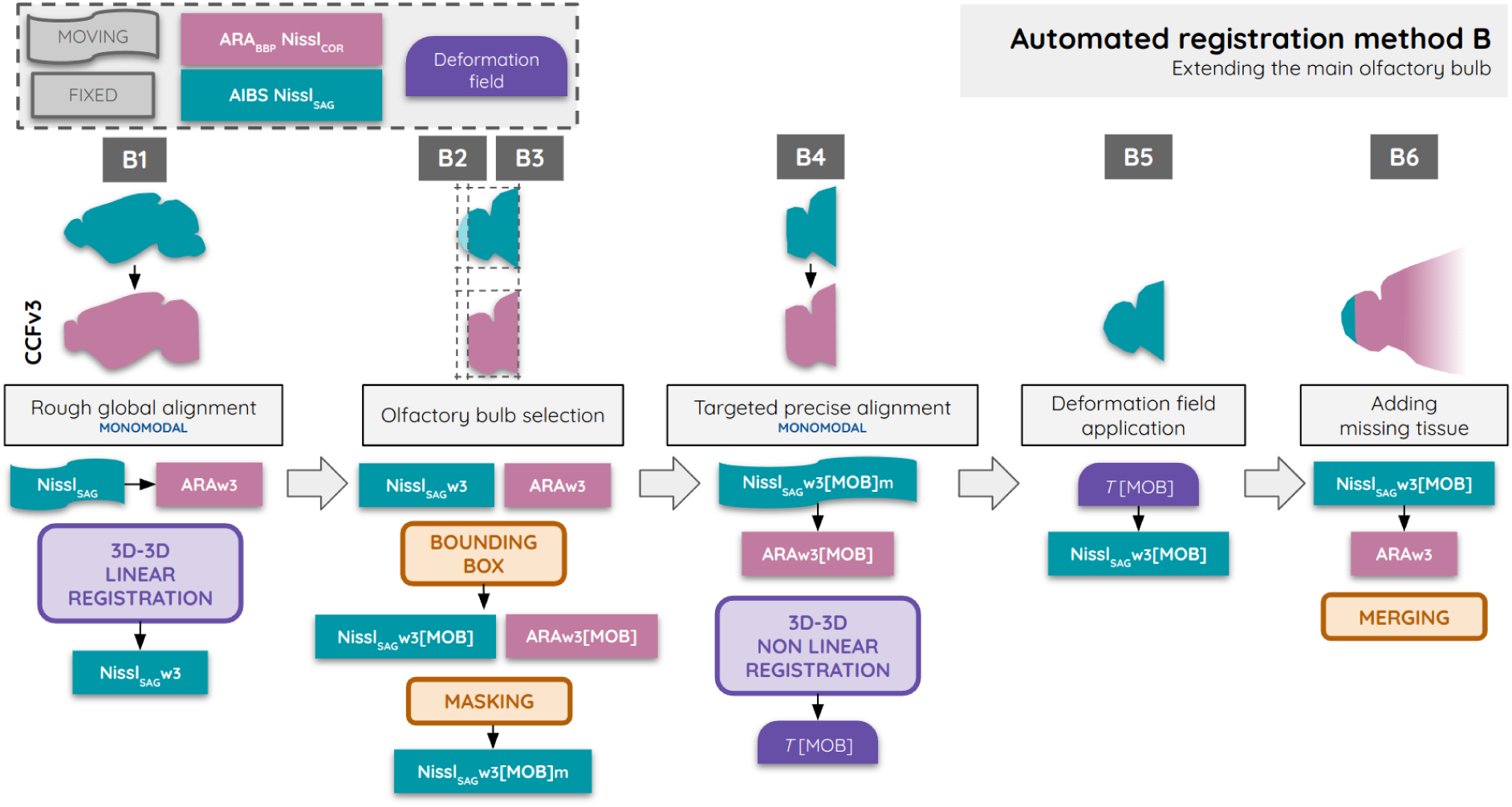
The six-step automated pipeline for adding the missing tissue from the AIBS Nissl_SAG_ to the ARA_BBP_ Nissl_COR_ in the CCFv3 olfactory bulb. The six steps for the ARM_B_ method are ranging from B1 to B6.

**FIGURE 6.**
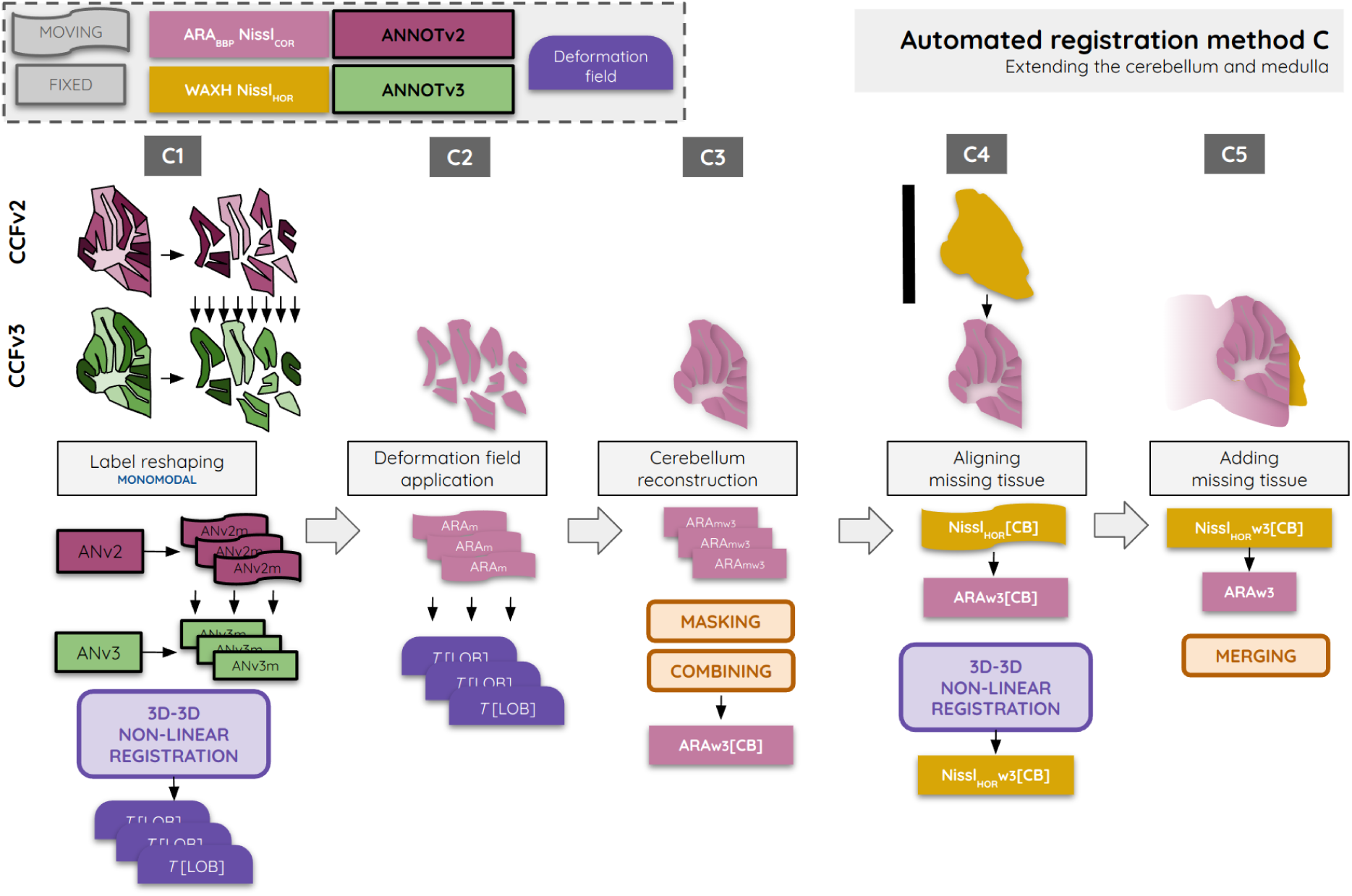
The five-steps automated pipeline for adding the missing layers in the ANNOTv3 as well as the missing tissue from the WAXH Nissl_HOR_ to the ARA_BBP_ Nissl_COR_ in the CCFv3 cerebellum. The five steps for the ARM_c_ method are ranging from C1 to C5.

We made use of freely available registration algorithms: Advanced Normalisation Tools (ANTs; Avants et al., 2008) and NiftyReg (Modat et al., 2014). These tools have demonstrated their efficiency for registering mono- and multimodal biomedical 2D and 3D datasets in various applications (Niedworok et al., 2016; Murakami et al., 2018; Nazib et al., 2018; Balakrishnan et al., 2019; Mano et al., 2020; Borovec et al., 2020; Iglesias et al., 2023). The ANTs algorithm is well-suited for precisely aligning small datasets, such as 2D histological slices, using nonlinear registration (Goubran et al., 2019; Krepl et al., 2021), especially its *symmetric image normalization* (SyN) transformation. However, it faces greater challenges when applied to large volumes at high resolutions (Johnson et al., 2023). Nonlinear 3D Nifty registration (NiftyRegF3D) registration has proven to be powerful in multimodal registration, since it is based on the normalized mutual information (NMI) similarity metric (Modat et al., 2014). We also used the visualization and segmentation software ITK-snap (Yushkevich et al., 2006) for assessing the resulting images.

Building on the CCFv3a_BBP_, we aligned 734 Nissl-stained mouse brains to the ARAva_BBP_ Nissl_COR_. From these sections we generated an average Nissl-stained template in the CCFv3a_BBP_. We produced a refined cell distribution placement within the atlas of the entire mouse brain given this new dataset (Rodarie et al., 2022). Furthermore, we merged the barrel column annotation produced by recent research (Bolaños-Puchet et al., 2024) in the cortex as well as the spinal cord model (Kuras, 2023) to our entire mouse brain CCFv3a_BBP_ annotation, thus producing a more complete annotation of the mouse central nervous system CCFv3c_BBP_ derived from the CCFv3. Volume concatenation and merging were used to produce such a refined and extended version.

The presented methods were conducted at 25 µm voxel isotropic resolution. The estimated deformation fields were then upsampled to 10 µm voxel isotropic resolution and applied to the raw Nissl-stained data at the same resolution as well. The resulting extended annotation, along with the granular and molecular layers across all lobules of the cerebellum, was upsampled to an isotropic resolution of 10 µm and smoothed.

Codes for the ARM_A_ method embed a high performance computing infrastructure for decreasing the computation time. This method was run using Blue Brain 5, which is an heterogeneous supercomputer with resources adapted to the scientific requirements of the project. In particular, we used 40 nodes from the cluster in which each node features two Intel Xeon Gold 6248 CPUs (*i.e.*, 40 core per node), 384 GB of memory, and are interconnected through Infiniband EDR. In total, our evaluations utilized 1600 processes distributed using the multiprocessing Python library (https://docs.python.org/fr/3/library/multiprocessing.html).

#### 1. Preprocessing

The WAXH Nissl_HOR_ was resampled at 25 µm and reconstructed in 3D using an adjacent linear registration technique (involves transformations that maintain straight lines) inspired by prior research (Ourselin et al., 2001). The latest ANNOTv3 provided on the AIBS platform was downsampled to an isotropic resolution of 25 µm using an in-house algorithm (see **Supplementary Material S3.1** for more details). Next, we removed some isolated and discontinuous voxels to improve discontinuity and jaggedness, located at the extreme borders of the 3D ANNOTv3, beyond the brain annotation volume itself (see **Supplementary Material S3.2**). The ANNOTv3 is not symmetrical; part of the dor*s*al tegmental decussation region, including 433 voxels, is present on the left hemisphere only (see **Supplementary Material S3.2**). As data are considered to be symmetric given the inter-hemispheric plane, we worked on a single hemisphere, and duplicated it at the end. We chose the right hemisphere for which the ANNOTv2 is better aligned to the ARA Nissl_COR_ (Krepl et al., 2021; Rodarie et al., 2022). Further, a portion of the main olfactory bulb region was approximated by inheriting the glomerular layer from its child region, as this layer was not present in the latest ANNOTv3 version (see **Supplementary Material S3.2**).

#### 2. Registering the ARA Nissl tissue in the CCFv3

For accurate registration of the ARA Nissl_COR_ within the CCFv3, resulting into ARA_BBP_Nissl_COR_, we developed the ARM_A_ based on the annotation parcellation (**Figure 2**). The idea is to restrict the registration on subregions to a comparable ontology. Although both annotations rely on the same hierarchy file, some regions exist in the ANNOTv2 that are not included in the ANNOTv3, and vice versa (see **Supplementary Material S4**). As those disparities are notably pronounced in the white matter regions – making them difficult to compare – our registration was focused on gray matter regions. Two ontology levels were defined: (1) the first is composed of all leaf regions (the term leaf represents the deepest level of the ontology) that are common in the two annotations, and (2) the second one is defined as an intermediate level of hierarchy (called the parent), where there is no common leaf labels between the annotations (see **Supplementary Material S4**). This second, intermediate ontology level is defined as being one of the first common parents between two annotations. In this context “common” refers to regions for which a non-zero set of voxels is defined with the same label in both annotation volumes. This approach ensures good precision of registration at the deepest ontology level (leaf) while concurrently filling in gaps where tissue is missing within a higher hierarchy level (parent). The objective of image registration is to find the optimal spatial transformation that aligns the *moving* image, on which the transformation is estimated and applied, with the *fixed* image (reference), ensuring that corresponding points in both images match as accurately as possible.

The following steps were serially applied in 3D for the ARM_A_ method (**Figure 4**):

**<u>Step A1</u>**: A monomodal linear, then nonlinear registrations (SyN) were performed between the ANNOTv2 (moving) and the ANNOTv3 (fixed) annotations using ANTspy (Krepl et al., 2021) as an initialization for making both annotations more comparable. The resulting transformation was applied to the ARA Nissl_COR_ with linear interpolation.

**Step A2**: The ARA Nissl_COR_ was subdivided into numerous subregions using the ANNOTv2 masks at the leaf and parent ontology levels independently, similarly for the STPT average template volume, where the ANNOTv3 was used.

**Step A3**: A linear, then nonlinear multimodal registrations were applied independently between each masked leaf and parent regions in the ARA Nissl_COR_ (moving) and in the STPT average template (fixed) using NiftyregF3D.

**Step A4**: The registered whole-brain tissue was reconstructed from Step A3, combining all the registered masked voxels from the leaf ontology and filling holes with the registered masked tissue from the parent region ontology.

**Step A5**: Monomodal nonlinear registration was applied at the whole-brain scale between the raw ARA Nissl_COR_ (moving) and the reconstructed ARA Nissl_COR_ from Step A4 (fixed) using NiftyRegF3D.

We used the default mode of the ANTs registration algorithm: initialization with affine registration (involving translation, rotation, scaling, and shearing), and then we applied the nonlinear SyN coupled to intersection over union similarity metric. The default mode of the NiftyRegF3D registration was also used, which is a free-form deformation based on the NMI similarity metric (NiftyRefF3D).

We used the NMI similarity metric to ensure better multimodal correspondence between the ARA Nissl_COR_ and the STPT average template (**Equation 1**). NMI calculation was performed: (I) at the whole-brain scale after the final step, (II) at the level of each registered subregion to ensure the homogeneity of the registering improvement across all subregions, and (III) at the level of individual slices, considering the volume as a succession of slices in the three conventional incidences (coronal, sagittal, and horizontal). In that manner, we provide a comprehensive depiction of whether the similarity between the two distinct anatomical volumes increased from these various perspectives. NMI is defined such as:

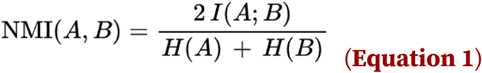

where *I*(*A*; *B*) is the mutual information between images *A* and *B*, *H*(*A*) is the entropy of image *A*, and *H*(*B*) is the entropy of image *B*.

We also used an independent metric to quantitatively assess whether the resulting aligned ARA_BBP_ Nissl_COR_ was better fitting the ANNOTv3. First, we defined a set of 29 fiducial anatomical points of interest in the ANNOTv3 clearly identifiable in the tissue based on good practices given in a previous study (Bjerke et al., 2019). These points were selected because they are distributed throughout the entire brain and are located at the boundaries between different regions. (see **Supplementary Material S6.1**). They also concern seven main regions of interest: olfactory areas, cerebellum, hippocampus, striatum, brainstem, thalamus, and cortex. Given this set of reference points readily identifiable, we tasked five operators with various profiles to identify their positions in the volume before (ARA Nissl_COR_) and after (ARA_BBP_ Nissl_COR_) registration (see **Figure 7K**). We evaluated and compared the target registration error (TRE) in calculating the Euclidean distance between each point defined by an operator and the reference set of points in the ANNOTv3 both before and after registration in the 3D volume (**Equation 2**), defined as:

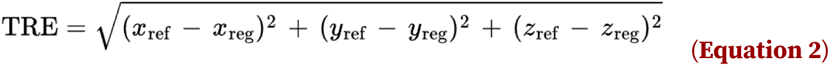

**FIGURE 7.**
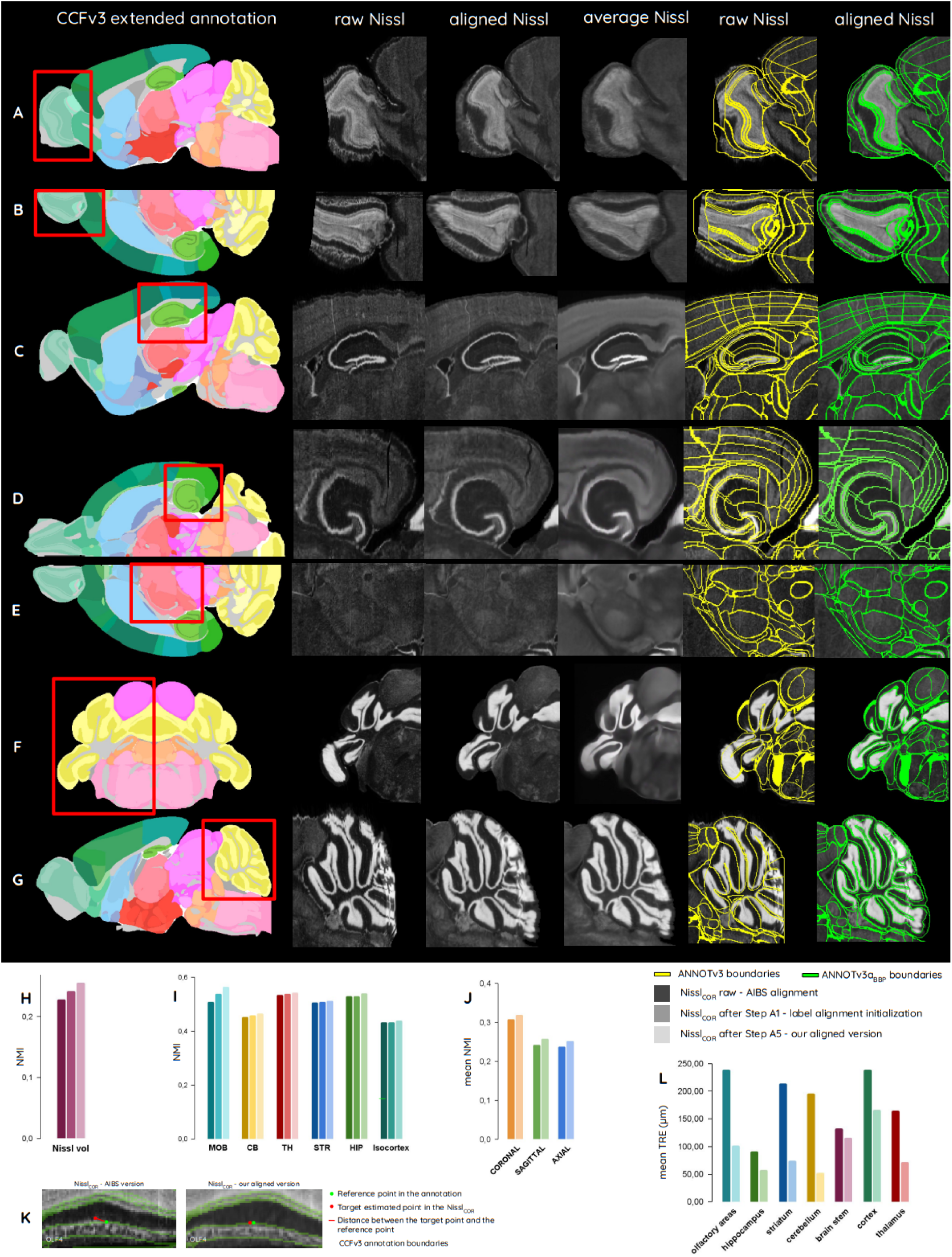
Aligning and extending the ARA Nissl_COR_ in the CCFv3 . (A-G) Sub-parts of the ARA Nissl_COR_ before (raw) and ARA_BBP_ Nissl_COR_ after (aligned) it was aligned in the CCFv3, as well as the ANNOTv3 boundaries raw (yellow) and extended ANNOTv3a_BBP_ (green); (H-J) NMI between the ARA Nissl_COR_ and the STPT average template (H) at whole-brain scale, (I) for six main groups of regions, and (J) for all slices in the three conventional incidences. (K-L) TRE calculated between the identified target points in the anatomy and the reference point in the annotation before and after alignment (K) for the example of point OLF4, and (L) for all the 26 points in main regions of interest (see **Supplementary Material S7.1** for more details).

where *x*_ref_, *y*_ref_, and *z*_ref_ are the coordinates of the fiducial point in the reference image (ANNOTv3), and *x*_reg_, *y*_reg_, and *z*_reg_ are the coordinates estimated in the ARA Nissl_COR_ before and ARA_BBP_ Nissl_COR_ after registration (Bookstein, 1989).

#### 3. Extending the rostral part of the Nissl brain

The AIBS Nissl_SAG_ was used to expand the ARA_BBP_ Nissl_COR_ reference, given that the entire olfactory bulb is available there. In addition, both datasets are Nissl-stained, making monomodal registration possible. Borders filled with zero values were added to the ARA_BBP_ Nissl_COR_ to fit the AIBS Nissl_SAG_ dimensions.

The following ARM_B_ method was developed to automatically incorporate missing tissue in the main olfactory bulb (here using the MOB acronym) from the AIBS Nissl_SAG_ into the ARA_BBP_ Nissl_COR_ (**Figure 5**):

**Step B1**: The AIBS Nissl_SAG_ (moving) was registered in 3D to the ARA_BBP_ Nissl_COR_ (fixed) using rigid then affine registration with the ANTs algorithm.

**Step B2**: Both the ARA_BBP_ Nissl_COR_ and the AIBS Nissl_SAG_ were cropped, corresponding to the bounding box for the olfactory bulb, resulting in the ARA_BBP_ Nissl_COR_[MOB] and the AIBS Nissl_SAG_[MOB].

**Step B3**: The AIBS Nissl_SAG_[MOB] was masked using the ANNOTv3 to hide the missing part of the olfactory bulb and match the ARA_BBP_ Nissl_COR_[MOB] tissue covering.

**Step B4**: The masked AIBS Nissl_SAG_[MOB] (moving) from Step B3 was nonlinearly (SyNOnly) registered to the ARA_BBP_ Nissl_COR_[MOB] (fixed) using ANTs algorithm, resulting in a transformation *T*[MOB].

**Step B5**: The transformation *T*[MOB] was applied to all voxels from the non-masked AIBS Nissl_SAG_[MOB] from Step B2.

**<u>Step B6</u>**: After a normalization of the AIBS Nissl_SAG_[MOB] using histogram matching with the ARA_BBP_ Nissl_COR_[MOB], all voxels from the nonlinearly registered AIBS Nissl_SAG_[MOB] from Step B5 were added to the ARA_BBP_ Nissl_COR_[MOB]. A 3-voxel thickness coverage along the rostro-caudal axis was used to average intensities around the junction, ensuring a smooth transition between the two brain parts.

The resulting main olfactory bulb Nissl-stained tissue extension was assessed by experts.

#### 4. Extending the caudal part of the Nissl brain

The WAXH Nissl_HOR_ was used to expand the ARA_BBP_ Nissl_COR_ reference, given that the entire cerebellum is represented in the dataset. Sixteen lobules of the cerebellum were concerned by the splitting into two layers which are the granular and molecular layers (see **Supplementary Material S8**): lingula (I), lobule (II), lobule (III), lobules (IV-V), declive (VI), folium-tuber vermis (VII), pyramus (VIII), uvula (IX), nodulus (X), simple lobule, crus 1, crus 2, paramedian lobule, copula pyramidis, paraflocculus, and flocculus.

The ARM_C_ method was used as follows (**Figure 6**):

**<u>Step C1</u>**: Each of the sixteen masked lobules was registered independently between the ANNOTv2 (moving) to the ANNOTv3 (fixed) with linear (affine), then nonlinear ANTs *aggressive symmetric normalization* (SyNAggro) algorithm coupled to the *demons* similarity metric (Avants et al., 2008).

**Step C2**: The sixteen 3D transformations estimated in Step C1 were applied to the sixteen masked ARA_BBP_ Nissl_COR_ lobules to align them within the CCFv3.

**Step C3**: The registered lobules were masked and merged in the reconstructed whole Nissl_COR_ volume from Step A4, generating a complete ARA_BBP_ Nissl_COR_ in the CCFv3, now including cerebellum in Step A5.

**Step C4**: The WAXH Nissl_HOR_ (moving) was registered to the ARA_BBP_ Nissl_COR_ (fixed) from Step C3 applying the same method used for the olfactory bulb in Steps B1-B4, but with a bounding box around the cerebellum.

**<u>Step C5</u>**: Since the WAXH Nissl_HOR_ was aligned in the CCFv3, it became possible to integrate missing data from the WAXH Nissl_HOR_ in the caudal part of the cerebellum with the ARA_BBP_ Nissl_COR_. We achieved this by applying the same steps to the WAXH Nissl_HOR_ as to the olfactory bulb (Step B5-B6) but with a bounding box around the cerebellum.

The medulla in the WAXH Nissl_HOR_ were integrated into the ARA_BBP_ Nissl_COR_ using the same method as applied to the olfactory bulb. This process is summarized in Steps C4 to C5.

The resulting cerebellum and medulla Nissl-stained tissue extensions were also assessed by experts.

#### 5. Filling the missing annotations into the ANNOTv3a_BBP_ version

After digitally reconstructing the entire Nissl olfactory bulb, cerebellum and medulla, it became possible to identify their layers using a semi-automated method in continuity with those defined in the ANNOTv3 to produce ANNOTv3a_BBP_ (see **Supplementary Material S3.4**). These annotation extensions and refinement concern 5 layers in the olfactory bulb (granular, inner plexiform, mitral, outer plexiform, and glomerular layers), 32 layers in the cerebellum (the granular and molecular layers in each of the 16 lobules listed in section 4), and one single layer in the medulla. In addition, the arbor vitae region from the fiber tracts was slightly manually extended in the uvula (IX) lobule to fit the extended version of the brain.

The semi-automated method we used for identifying all new layers in the ARAva_BBP_Nissl_COR_ anatomical data (see **Supplementary Material S3.4**) utilized different image processing techniques, among which binary Otsu thresholding (Otsu, 1975) was selected. This technique was especially successful in separating the denser layers from the other in each lobule of the cerebellum, independently. The granular layer was assigned to the highest intensity voxels (corresponding to high cell densities) and the molecular label to the low intensity (corresponding to low cell densities) according to the corresponding 32 leaf regions given in the hierarchy file (Šišková et al., 2013; Hayashi et al. 2017).

To assess the result from the semi-automated method (ANNOTv3a_BBP_), we compared it to the annotation file produced by experts manually (ANNOTv3a_m_). The procedure to obtain this ANNOTv3a_m_ is described in **Supplementary Material S3.3** (L’Yvonnet et al., 2024; Colnot et al., 2024). We did visual assessment in the three incidences using ITK-snap visualization software, and calculated Dice score (Dice, 1945) for each label as a quantitative analysis, taking ANNOTv3a_m_ as the reference (A) and ANNOTv3a_BBP_ as a target (B) (see **Supplementary Material S3.5**). Dice score (**Equation 3**) is defined as:

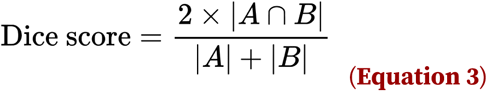

where |A⋂B| is the number of voxels that are common between the reference and target labels, |A| the number of voxels in the reference label A, and |B| the number of voxels in the target label B. This score ranges from 0 (no overlap) to 1 (perfect overlap).

The cerebellar Purkinje cells, precisely located at the junction between the granular and molecular layer of each cerebellar lobule, are defined by having a soma diameter ranging from 20 µm to 25 µm (Kim et al., 2011). Moreover, the thickness of the cerebellar Purkinje layer is comprised of a single layer of Purkinje cell somas (Šišková et al., 2013). A 25 µm voxel resolution is insufficient for delineating this region, as a curved single voxel layer will have discontinuities. However, as the ANNOTv3 version was also produced at 10 µm of resolution, it became possible to automatically generate a two 10 µm - voxel thin layer representing Purkinje layers at the exact boundary between the granular and molecular layer (one voxel in each layer) using image processing. This was a good compromise given the sampling conditions.

#### 6. Creating an average Nissl template

While we believe that a single Nissl-stained reference built from complementary brains (ARA_BBP_ Nissl_COR_, AIBS_BBP_ Nissl_SAG_, and WAXH_BBP_ Nissl_HOR_) may be more suitable for registration purposes, for some use cases it would be advantageous to additionally construct a population average Nissl template from many animals. For instance, when using Nissl intensity as an estimate of cell density (Rodarie et al., 2022), a population average of Nissl data representing many individuals would provide a more accurate approximation. To construct this, we downloaded 86,901 coronal Nissl-stained slices from 734 post-natal day 56 C57BL/6J mouse brains from the AIBS *in situ* hybridization data portal (Lein et al., 2007; Ng et al., 2007). We then used the DeepSlice registration algorithm to approximate the position of the sections in the CCFv3a_BBP_ (Carey et al., 2023). Although the AIBS provides sagittal sections, we did not use them because they cover only one hemisphere, and the current DeepSlice algorithm operates exclusively with coronal slices. This was followed by manual inspection of each dataset to correct for misalignments of angle and position, which were generally small. We then nonlinearly registered slices to their exact corresponding planes in the ARAva_BBP_ Nissl_COR_ using the SyN transformation from ANTspy registration library. We then placed the section values in their corresponding voxel in a 10 µm isotropic resolution volume in the CCFv3. We repeated these for each Nissl dataset summing the values together when two Nissl sections from different datasets intersected. In addition, we placed into this volume the intensities from the ARAv3a_BBP_ Nissl_COR_, AIBS_BBP_ Nissl_SAG_, and WAXH_BBP_ Nissl_HOR_, first averaging them together with the existing values. Since each Nissl dataset only covered a subset of the voxels, each voxel contained the sum of a variable number of datapoints. Then to correct for this, we calculated how many Nissl sections had intersected with each voxel. We then divided each voxel by the number of sections which intersected with it (in addition to the ARAv3a_BBP_ Nissl_COR_ volume). This produced a volume where each voxel contained the average Nissl intensity for that point from the datasets which intersected it.

## RESULTS

The ARA Nissl_COR_ was aligned in the CCFv3 and extended using the AIBS Nissl_SAG_ for the main olfactory bulb and the WAXH Nissl_HOR_ for the cerebellum and the medulla. We demonstrate in this section how the ARA_BBP_ Nissl_COR_ is better aligned in the CCFv3 than the original. Furthermore, we present the new extended ANNOTv3a_BBP_ version incorporating the granular and molecular layers in each lobule of the cerebellum. We also provide an augmented annotation covering the central nervous system directly using the ANNOTv3a_BBP_ we produced, in addition to computing the cell density in each region using our new ARAva_BBP_ Nissl_COR_ version.

### 1. Aligning and extending the ARA Nissl_COR_ in the CCFv3

We used image subsections of the ARA_BBP_ Nissl_COR_ in different incidences (**Figure 7A-G**) to qualitatively assess whether the matching between the anatomical volume and the extended ANNOTv3a_BBP_ has improved (see **Supplementary Material S7.1** for more details). To evaluate the similarity between the ARA Nissl_COR_ and the STPT average template before and after registration, we compared the NMI score at different stages of our pipeline: (1) before registration, (2) after Step A1, and (3) after Step A5. NMI scores were calculated (A) at the whole-brain scale (**Figure 7H**), as well as (B) at every level of the hierarchy (**Figure 7I**), and (C) for each slice in the three conventional incidences (**Figure 7J**). See **Supplementary Material S5** for more details. For (B), we presented the NMI scores only for the main brain regions, *i.e.* the main olfactory bulb, cerebellum, thalamus, striatum, hippocampal region, and isocortex. We also provided an example illustrating the TRE comparison for a given point in the list of fiducial reference points of interest (**Figure 7K**). Plots for the average TRE among the 26 points evaluated by five operators and divided into 6 regions of interest are presented in **Figure 7K**, see **Supplementary Material S6.2** for more details.

At the whole-brain scale, NMI scores between the STPT average template and the ARA_BBP_ Nissl_COR_ after Step A5 increased by 12% compared to the ARA Nissl_COR_, half of it being brought by the intermediate Step A1 (**Figure 7H**). For most regions, the alignment initialization (Step A1) was bringing almost half of the similarity improvements, except for the isocortex, where this step is responsible for only a 15% increase. The isocortex showed a relatively slight TRE decrease of 72 µm (**Figure 7L**). Although difficult to identify visually in the ARA_BBP_ Nissl_COR_, the cortical regions alignment improved by a small margin. Each of the main brain regions presented a similarity increase between 1% and 3% whereas the main olfactory bulb’s NMI score increased by 12% (**Figure 7I**). Qualitative analysis of the labels on the ARA Nissl_COR_ and ARA_BBP_ Nissl_COR_ in **Figure 7A-B** assessed a significant improvement in the poorly aligned regions. For instance, even the innermost layer (MOB, granule layer) extended beyond the MOB, outer plexiform layer in the raw ARA Nissl_COR_ (**Figure 7A**). In the improved version, each distinct layer has its own texture and contrast with its adjacent layers that are consistently matching the ANNOTv3 boundaries in the aligned version. In addition, TRE between points from that region presented one of the highest decrease of 137 µm on average (**Figure 7L**), also emphasizing the improvement in the alignment. Similar improvement was measured in the cerebellum. Qualitative analysis in **Figure 7F-G** showed some substantial tissue displacement, in particular for the most lateral lobule (paraflocculus) and the most rostral lobule (lobule II). TRE reduction for this region reports the best improvement among all regions, resulting on average in a 144 µm decrease after alignment (**Figure 7L**). However, some native artifacts are still present in the aligned version ARA_BBP_ Nissl_COR_ for that particular region, *e.g.* the vertical distortion of the tissue on the most caudal part of the region in **Figure 7G**. Note that this artifact is slightly smoothed by the alignment process. Consequently, the entire cerebellum has become less distorted than the raw version. The striatum also improved its positioning in the ANNOTv3 as it showed a significant TRE decrease of 140 µm on average (**Figure 7L**). With regard to the hippocampal region, only a slight improvement was observed, before the outermost a 2% increase in NMI, and an average decrease of 33 µm in TRE (**Figure 7I,L**). Slight alignment improvements were measured in the dentate gyrus which is visible from the sagittal view (**Figure 7C**) also reducing edge artifacts through the reduction of the distortion effect caused by the coronal tissue slicing. From a horizontal perspective, significant alignment improvements were observed for this thin region (**Figure 7D**).

Focusing on areas with higher contrasts, we can assess at least a slight and at best major improvement of the matching between the ARA_BBP_ Nissl_COR_ and the ANNOTv3 compared to the raw ARA Nissl_COR_ in regions under review (**Figure 7A-G**). Results regarding the slice-to-slice registration between the ARA Nissl_COR_ and the STPT average template before and after alignment showed NMI increase. The similarity with the STPT average template slices on average increased by 4% in the coronal incidence and by 6% in the sagittal and horizontal incidences (**Figure 7J**). Quantitative validations came to the same conclusion, both for the analysis of similarity, which increased overall in each region (**Figure 7H-J**), and for the TRE, which decreased on average across the surveyed regions (**Figure 7L**).

The tissues added to extend and complete the olfactory bulb and cerebellum integrated well with the continuity of the surrounding regions (**Figure 7A-B,G**). The extended main rostral part of the main olfactory bulb nevertheless remained blurred, making the junction between the raw and the added tissue visible (**Figure 7A-B**). Continuous connection and compatibility with the caudal part of the raw ARA Nissl_COR_ was achieved despite high distortion (**Figure 7G**).

Results for additional alignments of the AIBS Nissl_AXI_ and the WAXH Nissl_HOR_ in the CCFv3a_BBP_ are presented in **Supplementary Material S7.2**.

### 2. Producing the extended annotation of the central nervous system and the corresponding cell density in each region

The annotation brain volume changed between the different versions from the AIBS to ours: from 499.9 mm^3^ for the CCFv2 to 504.9 mm^3^ for the CCFv3, and finally to 511.7 mm^3^ for the CCFv3a_BBP_ (**Figure 2A-C**). The volume increased by 1.35% between the latest version from the CCFv3 and CCFv3a_BBP_, with the new version being closer to literature volume for the adult mouse brain, 508.9 mm^3^ (Badea et al., 2007). The new version incorporates an addition of 0.48 mm^3^ in the main olfactory bulb (2.6% of its total region), 2.52 mm^3^ in the cerebellum (4.5% of its total region), and 3.83 mm^3^ in the medulla (11.0% of its total region), see **Supplementary Material S9** for more details. The enlarged brain parts led to an increase of nearly 1 mm (950 µm) along the rostro-caudal axis, out of which 350 µm was dedicated to the main olfactory bulb, and 600 µm to cerebellum and medulla. In the main olfactory bulb, a larger area was affected by the extension, as part of the most rostral ANNOTv3a_BBP_ was modified to incorporate its layers (**Supplementary Material S8**). The total mouse brain length along the rostro-caudal axis increased from 13.2 mm to 14.1 mm, becoming closer to literature length, around 15 mm (Badea et al., 2007). All the new annotations incorporated in the ANNOTv3a_BBP_ benefited from qualitative and quantitative assessment using the expert annotation ANNOTv3a_m_. In particular, high dice scores in all regions (between 0.94 and 0.99) confirmed the close correspondence between those regions in ANNOTv3a_BBP_ and the expert annotation (see **Supplementary Material S3.5**).

Utilizing the DeepAtlas suite (https://github.com/BlueBrain/Deep-Atlas), we registered specific markers from the AIBS *in situ* hybridization coronal datasets to our extended atlas version CCFv3a_BBP_, using the extended annotation ANNOTv3a_BBP_ as well as the extended ARAva_BBP_ Nissl_COR_ (**Supplementary Material S10**). Most of the aligned slices fit well the ANNOTv3a_BBP_ boundaries, allowing the distribution of the markers to be used for quantitative analysis. We submitted the aligned markers with the extended ARAva_BBP_ Nissl_COR_ in the Blue Brain atlas pipeline (https://blue-brain-atlas-pipeline.kcpdev.bbp.epfl.ch/atlas-pipeline/) to produce cell densities for cell type compositions across the entire mouse brain. From these cell densities, we could determine the coordinates and labels of all cells in the CCFv3a_BBP_. A simulated Nissl-stained volume was produced from this data, representing each soma with a sphere (**Supplementary Material S11**). A simulated neuron atlas was then produced in the CCFv3a_BBP_ (**Figure 6E, Supplementary Material S11**).

In the main olfactory bulb (**Figure 2F**) it is possible to distinguish the different layers and their continuity in the entire region. Furthermore, layers in the extended part of the main olfactory bulb aligned well with the raw ARA Nissl_COR_ cell distribution (**Figure 7A-B**). A similar sectioned view of the cerebellum in **Figure 2G** made it possible to clearly distinguish the continuous boundaries between the granular and the molecular layer. The external layer of the extended part of the cerebellum annotation plus the boundary between the granular and molecular also fit well with the corresponding Nissl-stained areas in **Figure 7F-G**. The annotation boundaries were unaffected and preserved their smoothness despite the distortion artifact that remains in the caudal part of the cerebellum in **Figure 7G** which was not fully corrected by the alignment process.

By combining more than 86,000 registered coronal Nissl-stained slices from 734 mouse brains, we produced an averaged Nissl-stained volume aligned in the CCFv3a_BBP_ from our aligned version ARAva_BBP_ Nissl_COR_ (**Figure 7A-G**; see **Supplementary Material S7.3**). The first initialisation of this data was the average of ARAva_BBP_ Nissl_COR_, AIBS_BBP_ Nissl_SAg_, and WAXH_BBP_ Nissl_HOR_ (see **Supplementary Material S7.2**). This averaged Nissl-stained volume exhibits enhanced smoothness, with certain structures, such as the ventral posteromedial nucleus of the thalamus and the corpus callosum, emerging more clearly than in a single brain (**Figure 7D**), and the junction between the layers 4, 5, and 6 of the isocortex. Although some tissue damage is still visible, the data has significantly fewer artifacts. For instance, we eliminated the tissue tear in the lateral part of the isocortex close to the hippocampus (**Figure 7D**) .

To create a more complete mouse brain model, we first added the barrels annotation in the extended ANNOTv3a_BBP_ (**Figure 2D**). Then, we connected the spinal cord annotation and its corresponding cell densities to the caudal part of the brain. As a result, a continuous mouse central nervous system annotation (CCFv3c_BBP_) emerged without any cut in it: from the most rostral part of the main olfactory bulb to the very end of the spinal cord (**Figure 2D**).

All data produced here are freely available (https://doi.org/10.5281/zenodo.13640418). This includes every version (listed in **Table 1**, BBP section), plus the aligned AIBS_BBP_ Nissl_SAG_ and WAXH_BBP_ Nissl_HOR_ in the CCFv3a_BBP_, enabling usage in different contexts. The main version was uploaded to the BrainGlobe API (https://brainglobe.info/index.html), making the new atlas available on this interface (Blixhavn et al., 2024). Code related to the production of all the data is open source and can be downloaded at https://doi.org/10.5281/zenodo.13640418.

## DISCUSSION

We generated a new mouse brain atlas derived from the AIBS CCFv3, which includes the reconstruction and annotation of previously missing tissue in the rostral and caudal regions. Continuity and compatibility between the raw ARA Nissl_COR_ and the extended tissue was assessed. We also aligned the ARA Nissl_COR_ in the CCFv3 which significantly improved its fitting with the ANNOTv3. The molecular and granular layers were also added to the ANNOTv3a_BBP_, fitting the Nissl-stained anatomy in the CCFv3a_BBP_. This enabled production of a smooth averaged Nissl-stained volume, and a simulated Nissl-stained mouse brain volume where coordinates and labels of each cell are known.

### 1. Adding the missing tissue in the ARA_BBP_ Nissl

To incorporate the missing tissue in the Nissl-stained sectionsl, we had to meet both targets: use biological, original Nissl-stained data and correspond to the geometry given by the CCFv3. Rigid registration, our first step, was not sufficient for creating a correspondence between the AIBS Nissl_SAG_ or the WAXH Nissl_HOR_ and the ARA_BBP_ Nissl_COR_. We thus used affine registration (Step B1, Step C4) for making the volumes correspond despite their different sectioning artifacts. An affine registration was performed at the whole-brain scale to maintain all structural correspondences between volumes, including the continuity beyond the range of the ARA_BBP_ Nissl_COR_. This constraint made it possible to preserve the 3D consistency of the reconstructed anatomy in the CCFv3.

The new tissues added to the ARA_BBP_ Nissl_COR_ volume were successfully labeled: the main olfactory bulb, cerebellum, and medulla. The main olfactory bulb labels were subdivided into five layers and the cerebellum labels were further subdivided into lobules, and then into three layers corresponding to their respective leaf regions. The medulla did not benefit from such subdivision, as the WAXH Nissl_HOR_ did not provide sufficient information to identify its leaf regions in the added tissue. To achieve this level of detail, it will be necessary to assess such subdivisions under expert supervision integrating new data including specific markers in the future.

The three Nissl-stained volumes were registered in the same CCFv3 coordinates to complement each other and to reconstruct missing tissue. These additions provide a comprehensive coverage of the entire mouse brain, each volume constructed predominantly of horizontal, sagittal or coronal sections (see **Supplementary Material S7.2**). After the construction of the extended ARAva_BBP_ Nissl_COR_, several regions needed to be considered carefully. Depending on the viewing incidence, the tissue appeared blurred in the extended regions. Hence, for any slice-to-slice registration, we recommend selecting the volume cut in the same incidence in which was used for producing the experimental slices. This would result in an even better alignment by ignoring the distorted 3D reconstruction effect.

As the volume dimensions increased along the rostro-caudal axis of the brain, the zero origin coordinate along this axis was shifted. Since this would change the reference space coordinate initially defined by the AIBS, we chose to include a negative shift of 350 µm in the rostral direction. This way all coordinates from the original CCFv3 were preserved.

### 2. Adding the granular and molecular layers in the cerebellum

The ARM_A_ method we proposed for aligning the ARA Nissl_COR_ within the CCFv3 on a region-by-region basis was insufficient for accurately registering the cerebellum due to several factors. First, the absence of distinct labels differentiating the granular and molecular layers in the ANNOTv3 complicated the alignment process. The hierarchical level of cerebellar lobules in ANNOTv3 was not precise enough to segment the ARA Nissl_COR_ adequately, making accurate cerebellar region registration within the CCFv3 difficult. Second, the combination of inter-individual variability and averaging in the STPT average template resulted in a blurred anatomical separation between the granular and molecular layers, making these layers impossible to identify within the tissue. Lastly, the distorted nature of ANNOTv2 in this particularly folded region further added to the complexity of the registration process. Overall, a dedicated method was needed for the cerebellum (ARM_C_), involving the identification of the granular and molecular layers before running the last monomodal registration Step A5 (see **Figure 3**).

The cerebellar cortex is composed of lobules that are subdivided into molecular and the granular layers in the ANNOTv2, whereas in the ANNOTv3 lobules are not divided. Our method introduced clear layer distinction for each lobule in the ANNOTv3. Using lobule annotation registration, we could have just added the granular and molecular labels from the ANNOTv2 to the ANNOTv3 in the cerebellum. However, as those labels were distorted, we preferred to recreate them based on Nissl-stained data, where they are clearly identifiable. This also improved the 3D smoothness of the ANNOTv3. In our new annotation version, the layers appeared realistic due our data-driven method. We paid particular attention to preserving the labels of the lobules and the arbor vitae regions. This was done to maintain a shared reference coordinate system for the scientific community. We only incorporated the granular and molecular layers of the cerebellar lobules to the volume, leaving the rest unchanged.

Due to the higher cell density in the granular layer compared to the molecular layer, the boundary between these layers was clearly visible in single brain Nissl-stained volumes. However this characteristic became undetectable in the STPT average template, which is derived from the average of more than one thousand brains. This boundary may vary for each brain due to inter-individual variability in that region and potential artifacts that often occur on the outer edge of the tissue during extraction. Registering Nissl-stained lobules with the STPT average template lobules was difficult because the process often distorted either the granular or molecular layer to match the smooth, featureless STPT average template lobule. As a result, the lack of contrast rendered the STPT average template unsuitable for aligning the cerebellar Nissl-stained tissues. That is why we selected a more permissive and powerful registration algorithm for the lobules as we were constrained to only use the annotation files (ANNOTv2 and ANNOTv3). The SyNAggro transformation (fine-scale matching and more deformation) coupled to the demons similarity metric from ANTs was chosen. The task was brought to match two different empty shapes with no contrast inside (such as in Step A1). Since the transformation was computed for each voxel individually, the ARA Nissl_COR_ tissues also conform to this overall deformation.

The reconstructed WAXH Nissl_HOR_ was chosen as the best dataset for automatically identifying the granular layers throughout the entire cerebellum. In particular, the WAXH Nissl_HOR_ was smoother across all three incidences and exhibited fewer artifacts in certain lobules compared to the ARA Nissl_COR_. As a result of this selection, the granular layer annotation added to the CCFv3a_BBP_ is free from the artifacts present in the ARA Nissl_COR_. This implies that any monomodal registration applied to our extended ARAva_BBP_ Nissl_COR_ can be used to identify cerebellar layers in other Nissl-stained volumes.

### 3. Aligning the Nissl-stained tissue in the CCFv3

Registering the ANNOTv2, the ANNOTv3, the STPT average template, and the ARA Nissl_COR_ dataset in various ways presented several significant challenges. First, we attempted to register the ANNOTv2 to the ANNOTv3. A preliminary rough alignment at a comparable ontology level slightly enhanced the registration quality when applied to the ARA Nissl_COR_ volume despite the significant inherent constraints posed by the distortions and discontinuities in the ANNOTv2. This improvement, though modest, was valuable as an initialization step because it reduced distortions in the CCFv2. Second, our efforts to register the ARA Nissl_COR_ to the STPT average template revealed that current nonlinear registration algorithms are not powerful enough to effectively handle whole-brain scale multimodal registration. Achieving a global optimal registration is challenging due to the need to simultaneously optimize nonlinear 3D transformations across tens of millions of voxels, further complicated by variations in contrast and anatomical multimodality. The distinct modalities and dimensionalities (2D histology for the ARA Nissl_COR_ versus 3D serial two-photon tomography for the STPT average template), as well as the fact the ARA Nissl_COR_ is a single brain while the STPT average template is an average of 1,675 brains, further exacerbated the challenge. Third, the fundamentally different modalities of anatomical tissue versus descriptive labels interpreting regions, made direct comparison and alignment unfeasible, as the ARA Nissl_COR_ alone could not consistently establish clear region boundaries. Additionally, the lack of a suitable similarity metric for this hybrid registration problem highlighted the absence of adapted tools necessary for effective registration. All these challenges led us to propose a region-based registration method for registering one atlas with another.

We also tried to apply the exact same method at 10 µm isotropic resolution and assessed that similarities between the STPT average template and the ARA Nissl_COR_ improved. However, some regions presented poorer registration quality compared to the 25 µm version. Registering regions with an even higher number of voxels made the registration algorithm struggle to estimate a global transformation that maximizes coverage between highly different contrasts within images. At this resolution, differences in contrast were heightened and their variety increased, increasing the potential for misalignment. Moreover, the 10 µm region-by-region alignment process and its validation required a much longer computing time. Ultimately, we chose a resolution of 25 µm for our registration pipeline. Nevertheless, we produced the 10 µm isotropic resolution version by upsampling the deformation field rather than computing it from the native datasets. We assumed that the low amplitude of the deformations would ensure a smooth and coherent registration process. As a result, contrasts present in the native ARA Nissl_COR_ data were preserved and it was not blurred due to oversampling.

The AIBS annotation expertise made it possible to perform focused registration of the same anatomical tissue within the two volumes (region-to-region registration, Steps A2-A4). This method has the advantage of preserving the alignment process of the influence of contrasts in neighboring regions, ensuring that no point outside the concerned region can be registered from the moving space to the reference space. This method also ensured that no region was disadvantaged relative to another during the registration process regardless of their contrast differences. On one hand, when specific contrast was present in the same region across both anatomical volumes,, the registration process ensured its preservation. On the other hand, when there was no particular contrast *i.e.* the region was homogeneous, virtually no deformation was applied and the tissue was simply transferred from one modality to the other. The result of the registered main olfactory bulb well illustrates this layer-based alignment process.

It was also important to mask the registered data in order to save only the most useful signal and remove the rest of the distorted tissue, making subvolumes smooth (Steps A4). This approximation was based on the assumption that most of the tissue was well registered, enough for providing key features in the reconstructed volume in Step A4 for the final registration process in Step A5. Indeed, monomodal registration led to precise tissue alignment, which was not possible before. This last alignment step made it possible to retrieve 3D consistency without any tissue loss. The nonlinear registration made transformations quite permissive, but at the same time transformations were constrained by the regions themselves. Further, the independent registration in Step A5 resulted in realistic, smooth deformations of low amplitude while minimizing tissue deformation. This could explain why the NMI score increased by only 12%, the alignment resulting in many slight deformations among the whole 3D volume while preserving all tissue.

This region-based atlas alignment method can be applied to other atlases in similar contexts. By combining labels and anatomical tissues, our method addresses the multimodal registration issue, making a significant contribution to the field of multimodal atlas registration. Such a work was recently initiated with light sheet fluorescence microscopy and magnetic resonance imaging data (Perens et al., 2023). There is still work needed to integrate most reference atlases into a single, common coordinate framework, such as the Paxinos (Paxinos et al., 2019) or the Dorr (Dorr et al., 2008) atlases for instance.

All major brain regions reviewed showed a significant overall increase in the fitting of the ARA_BBP_ Nissl_COR_ with the CCFv3. However, it is important to acknowledge the limitations. The results were not uniform and varied from region to region. Quantitative results showed that the increase in NMI and decrease in TRE were less pronounced in the isocortex and hippocampus compared to regions like the cerebellum and main olfactory bulb. At this scale, the ARA_BBP_ Nissl_COR_ isocortex does not allow for the visual detection of all layers defined in it in the ANNOTv3a_BBP_, due to the lack of significant intensity differences. The alignment process only produced slight improvements in this region, due to poor contrast. This could explain the low improvement scores. Regarding the hippocampus, the most notable anatomical contrast specifically involves the dentate gyrus. Only precise parts of this very thin and dense region were misplaced in the ANNOTv3. The quantitative analysis does not fully capture the extent of alignment improvements in that region due to the limited number of evaluated points, which may account for the relatively minor global improvement calculated for the hippocampus. The main olfactory bulb was one of the most misaligned regions, which presented one of the highest alignment improvement scores with our version. The same situation applied to the cerebellum, where most lateral and caudal lobules were particularly misaligned. In practice, it was hard to preserve the 3D structure integrity of these regions, more specifically for the paraflocculus which stands apart from the rest of the brain. Both the main olfactory bulb and the cerebellum presented substantial differences in intensity (high contrast) and texture between their different layers. This made it possible to register each layer separately, retrieving this intensity and texture difference in the CCFv3. The final alignment stage (Step A5) enabled each layer to be placed in its proper position.

Similarity scores improved in all three conventional views following the alignment. It is interesting to note that the method resulted in only minor improvements in similarity scores in the coronal plane compared to the other two planes. This indicates that the alignment efforts have already been undertaken in the coronal plane (native cutting plane), compared to the sagittal and horizontal planes. In the end, our region-based alignment method could overcome the alignment difficulties between the ARA Nissl_COR_ with the STPT average template, in particular in sagittal and horizontal views. Consequently, jagged-edge artifacts were smoothed out in these two planes.

Observing more in depth, the NMI scores related to the main olfactory bulb similarity increased a lot more compared to the cerebellum after Step A5. We could deduce that improvement was not high in the cerebellum because of the lack of contrasts in the STPT average template compared to the ARA_BBP_ Nissl_COR_. Here we encountered a limitation of the similarity metric, which failed to accurately depict the intensity of similarity improvement in the cerebellar region due to multimodality. This is why we decided to evaluate the alignment of the ARA_BBP_ Nissl_COR_ with the CCFv3 using the TRE as an additional measure. To design the fiducial points of reference, we selected locations at the boundaries between regions. This approach was preferred over selecting points within regions because it is simpler and more effective in ensuring that they are easily identifiable in the 3D volume. Additionally, points placed within a region might not be as relevant for evaluating the alignment of the tissue within the annotation, as centers are less sensitive to the alignment accuracy than region boundaries.

The raw ARA Nissl_COR_ included some artifacts, in some lobules of the cerebellum and the caudal part of the isocortex. We made the choice not to correct such artifacts, which sometimes were a little spread by the registration process. In this manner, we preserved the ARA Nissl_COR_ volume in its original version and avoided intense manual processing on it. Especially, the caudal part of the cerebellum continues to suffer from a significant artifact that was present in the ARA Nissl_COR_. The arbor vitae region remains misaligned with the Nissl tissue. This misalignment is pronounced in the declive (VI) lobule. The ARA_BBP_ Nissl_COR_ version, while not perfect, is adequate for accurately aligning individual histological slices or volumes in CCFv3, serving as a crucial transitional dataset with improved precision compared to previous versions.

This paves the way for massive registration in the CCFv3a_BBP_ of gene expression databases such as the AIBS *in situ* hybridization data for instance (Lein et al., 2007; Ng et al., 2007). Most of the slices we used are well aligned, but misalignments were still found (see **Supplementary Material S10**). The registration algorithm was prone to errors, further compounded by the assumption that the plane was perfectly coronal, a condition that is rarely met in practice. Despite misalignments, precise quantitative analysis of gene expression in the CCFv3a_BBP_ was performed on that dataset. We used the averaged ARAva_BBP_ Nissl_COR_ for refining the cell distribution estimation. By realistically simulating the 3D ARAva_BBP_ Nissl staining in the CCFv3a_BBP_ in the BioExplorer software ©, we could navigate through the brain, to see each cell and its corresponding region. It is therefore possible to generate any cross-section and imagine any experimental design for simulating Nissl staining of any region in 3D (see **Video V1**).

The production of the average Nissl template in the CCFv3a_BBP_ increased the applicability of the new version of the atlas. The averaged Nissl-stained template became more compatible with other templates and could favor multimodal studies, in addition to being almost artifact-free. Three essential elements were required to produce it: (1) the DeepSlice tool for precisely identifying each slice position in the CCFv3a_BBP_, (2) the ANTs registration algorithm for producing massive nonlinear slice-to slice registration, and (3) the reference data in the CCFv3a_BBP_ to bring all data into the atlas, which is the ARAva_BBP_ Nissl_COR_ we produced. The smooth contrasts, combined with the clear highlighting of major structures and minimal visible artifacts, demonstrated that our alignment quality was sufficient for the registration algorithm to achieve accurate results. Indeed, the visible transitions between the raw ARA Nissl_COR_ and the extended part disappeared. Despite some artifacts, misaligned tissues, and the fact that the entire pipeline was run at an isotropic resolution of 25 µm, we have shown that our alignment is robust enough for the ARAva_BBP_ Nissl_COR_ to serve as a powerful tool for cross-space data transition at an isotropic resolution of 10 µm. The use of such smooth 3D data extends the capability to register sections in any plane or subvolume while preserving smooth reference data. This is particularly useful for the QuickNII tool (Puchades et al. 2019), as well as for registering 3D histology data produced by light-sheet fluorescence microscopy (Renier et al., 2016). Aligning all the sagittal Nissl-stained slices in our CCFv3a_BBP_ atlas would certainly refine this average Nissl template. It can facilitate the creation of a multimodal pool of anatomical data within the extended CCFv3a_BBP_, integrating recent research datasets from light sheet fluorescence imaging and magnetic resonance imaging 3D scans (Voigt et al., 2019; Perens et al., 2023) for instance.

The average Nissl template represents the average cellular density in the mouse brain, making it ideal for generating a simulated volume containing the position and type of each cell (based on its morphological-electrical property for the neurons), and the anatomical region to which it belongs (see **Supplementary Material S11**). This allows us to subsequently connect the simulated circuits in order to build and simulate the mouse central nervous system *in silico*. The digital twin of an entire Nissl-stained mouse brain is also valuable for neurobiologists. Having captured the exact position of any experimental slice during the cutting process on the microtome made it possible to directly generate its corresponding Nissl-stained slice without having to produce it, or even directly using DeepSlice without any preliminary information. This dataset offers the same information as a dataset obtained through cell soma counterstaining.

Connecting the spinal cord to the extended annotation made it possible to propose a comprehensive CCFv3c_BBP_ atlas covering the central nervous system from the most rostral point of the main olfactory bulb to the end of the spinal cord. By integrating complementary or more detailed annotation models, such as those for the spinal cord or barrel columns, we demonstrated that any contribution can be incorporated into a unified version of the atlas. Our goal has been to develop an atlas model aligned with the one proposed by AIBS, ensuring it is comprehensive, versatile, and openly accessible, thereby enabling others to contribute their insights and further improve it. All data will also be integrated on BrainGlobe to connect with other models and enhance visibility within the scientific community.

## CONCLUSION

Our work represents a significant contribution in the field of mouse brain atlases, as we have produced an extended and improved version of the mouse brain atlas derived from the AIBS CCFv3. By carefully addressing the limitations of incomplete brain coverage and alignment inaccuracies, we have reconstructed the missing tissue in both the rostral and caudal parts of the brain, while also incorporating layered differentiation in the cerebellum. Through our region-based automated and reproducible registration method, we have substantially improved the alignment of the Nissl-stained volume in the CCFv3. We bridged the gaps between the native truncated CCFv3 atlas of the mouse brain and both the aligned Nissl-stained volume as well as the mouse central nervous system atlas. This comprehensive atlas supports a broad spectrum of applications, from histological analyses to computational modeling.

## DATA AND CODE AVAILABILITY

All data and code produced in this study are freely available at https://doi.org/10.5281/zenodo.13640418. An additional file including codes in the format of a toolbox for producing is also available. Following the pipeline description in the paper will allow you to reproduce the data.

## AUTHOR CONTRIBUTIONS

H.M. and J.G.B. Funding Acquisition, Resources and Supervision. A.R. Supervision. D.K. Conceptualization, Supervision, and Project Administration. T.L. and E.C. Data curation, expert assessment, manual annotations for the main olfactory bulb and the cerebellum, respectively, and landmark identification. E.D. Methodology, Data curation, technical support and landmark identification. H.C. Data curation. C.V. Validation, Methodology, Data curation, scientific support and landmark identification. S.P. Conceptualization, Data Curation, Investigation, Formal analysis, Methodology, Validation, Visualization, landmark identification, and Writing - original draft. All authors prepared the manuscript Writing - Review & Editing.

## ACKNOWLEDGMENTS

This study was supported by funding to the Blue Brain Project, a research center of the École polytechnique fédérale de Lausanne (EPFL), from the Swiss government’s ETH Board of the Swiss Federal Institutes of Technology. This project/research has received funding from the European Union’s Research and Innovation Program Horizon Europe under Grant Agreement no. 101147319 (EBRAINS 2.0). Figures with 3D visualization as well as videos were produced by Cyrille Favreau using Blue Brain BioExplorer 1.8.0. We thank Karin Holm for editorial assistance.

## DECLARATION OF COMPETING INTEREST

The authors declare no competing interests.

## Notes

https://doi.org/10.5281/zenodo.13640418

### Competing Interest Statement

The authors have declared no competing interest.

