## Supplementary Material for "An extended and improved CCFv3 annotation and Nissl atlas of the entire mouse brain"

#### S1 Acronyms

**AIBS**: Allen Institute for Brain Science

**AIBS Nissl$_{SAG}$**: whole mouse brain from the AIBS sectioned in the sagittal incidence

**AIBS$_{BBP}$ Nissl$_{SAG}$**: AIBS sagittal Nissl-stained volume aligned in the CCFv3a$_{BBP}$ at BBP

**ANNOTv2:** CCFv2 annotation volume

**ANNOTv3:** CCFv3 annotation volume

**ANNOTv3a$_{BBP}$:** extended CCFv3 annotation produced at BBP covering the entire mouse brain and including new cerebellar layers

**ANNOTv3c$_{BBP}$:** extended CCFv3a$_{BBP}$ annotation produced at BBP covering the mouse central nervous system including spinal cord as well as barrel columns

**ANTS:** Advanced Normalisation Tools

**ARA Nissl$_{COR}$**: Allen Reference Atlas coronal Nissl-stained volume

**ARA$_{BBP}$ Nissl$_{COR}$:** ARA Nissl$_{COR}$ accuratelay aligned in the CCFv3 at BBP

**ARAva$_{BBP}$ Nissl$_{COR}$:** ARA Nissl$_{COR}$ accuratelay aligned and extended in the CCFv3 at BBP

**ARM$_A$:** automated registration method for aligning the ARA Nissl$_{COR}$ into the CCFv3

**ARM$_B$:** automated registration method for extending the main olfactory bulb Nissl

**ARM$_C$**: automated registration method for extending the cerebellum as well as the medulla Nissl, plus adding the missing cerebellar granular and molecular layers.

**BBP:** Blue Brain Project

**CCF**: Common Coordinates Framework

**CCFv2**: Common Coordinates Framework version 2

**CCFv3**: Common Coordinates Framework version 3

**CCFv3a$_{BBP}$**: Entire mouse brain atlas derived from CCFv3 produced at BBP

**CCFv3c$_{BBP}$**: Mouse central nervous system atlas derived from CCFv3 produced at BBP including spinal cord and barrel columns

**NiftyRegF3D:** non-linear 3D Nifty registration

**NMI:** normalized mutual information

**STPT**: serial two photon tomography

**SyN:** symmetric image normalization

**TRE:** target registration error

**WAXH Nissl$_{HOR}$**: whole mouse brain from the Waxholm space sectioned in the horizontal incidence

**WAXH$_{BBP}$ Nissl$_{HOR}$**: Waxholm horizontal Nissl-stained volume aligned in the CCFv3a$_{BBP}$ at BBP

### S2 Data location

All data are in open access and were downloaded from the following links:
© 2015 Allen Institute for Brain Science. Allen Brain Atlas API. Available from: brain-map.org/api/index.html

- Allen Mouse brain atlas (2011):
  - CCFv2 annotation (ANNOTv2): https://download.alleninstitute.org/informatics-archive/current-release/mouse_ccf/annotation/mouse_2011/)
  - coronal Nissl-stained volume (ARA $Nissl_{COR}$): https://download.alleninstitute.org/informatics-archive/current-release/mouse_ccf/ara_nissl/)
- Allen Mouse brain atlas (2022):
  - CCFv3 annotation 2022 (ANNOTv3): https://download.alleninstitute.org/informatics-archive/current-release/mouse_ccf/annotation/ccf_2022/
  - average template: https://download.alleninstitute.org/informatics-archive/current-release/mouse_ccf/average_template/
- hierarchy file: http://api.brain-map.org/api/v2/structure_graph_download/1.json
- sagittal Nissl-stained mouse brain volume (AIBS $Nissl_{SAG}$): https://mouse.brain-map.org/experiment/show/100042147
- horizontal Nissl-stained mouse brain from the Waxholm space (WAXH $Nissl_{HOR}$): https://www.nitrc.org/projects/incfwhsmouse

### S3 Pre- and post-processing on the CCFv3 latest version

#### S3.1 CCFv3 atlas downsampling

To create a 25 µm isotropic resolution version of the annotation volume, we used slicing to downsample the Allen Institute's 10 µm isotropic resolution volume. A set of procedures were applied to the 10 µm isotropic resolution.

First, we defined the scaling factor as 2.5 to reflect adjusting from 25 µm to 10 µm, which corresponds to reducing each dimension of the array by a factor of 2.5.

Next, we computed the new shape of the array as a tuple of integers, where each dimension is divided by the scaling factor and rounded down to the nearest integer.

Then, we used the `numpy.linspace()` function to generate a set of evenly spaced indices along each dimension of the array. The `linspace()` function takes three arguments: the start value (0), the stop value (dim-1), and the number of indices (new_dim). The resulting indices are floating-point numbers, so we convert them to integers using the `astype()` method.

Finally, we used the `numpy.ix_()` function to select a subarray of the original array using the indices. The `ix_()` function takes one or more indexing arrays as arguments and returns a tuple of arrays that can be used to index into the original array. We then stored the selected subarray in the new_atlas variable and print the shapes of the original and new arrays to verify that the new array has the desired shape of (528, 320, 456).

This downsampling step inherently introduces artifacts, as it either enlarges the relative size of brain regions smaller than a 25 $\mu m^3$ edge voxel or completely removes them from the annotation volume. If we look at the symmetric difference between the two annotation versions after slicing, only the regions field CA1 (382), field CA3 (463), retrosplenial area, dorsal part, layer 4 (545), and direct tectospinal pathway (1051) disappeared from the downsampled annotation volume. Note, that the original size (10 µm isotropic resolution) of these regions were only 6, 2, 2, and 4 voxels, respectively. While taking note, we deemed these changes acceptable, as the alternative would have entailed significantly enlarging the relative size of these small regions. Code is available upon request.

#### S3.2 Annotation corrections

In the original 10 µm isotropic resolution annotation volume from the Allen Institute, four isolated and discontinuous voxels, located at the extreme borders of the volume (most rostral coronal plane), were detected. We assumed these voxels were errors and removed them from the annotation file before downsampling. They were labeled as being field CA3, stratum lacunosum-moleculare (471). The coordinates of those four voxels are [1319, 799, 520], [1319, 799, 521], [1319, 799, 618], and [1319, 799, 619].

The latest version of the CCFv3 annotation is not symmetrical. In particular, the dorsal tegmental decussation (1060) contains 430 more voxels in the left hemisphere compared to the right hemisphere, all located in the most medial left sagittal plane.

Moreover, three more discontinuous points are only present in the left hemisphere, two from the main olfactory bulb, mitral layer (236) located at positions [245, 500, 524] and [245, 501, 524], and one from the Lateral septal nucleus, rostral (rostroventral) part (258) at position [548, 313, 521]. In this work, we focused only on the right hemisphere, eliminating all these points.

Part of the main olfactory bulb (507) label was re-labeled by an expert into the main olfactory bulb, glomerular layer (212) as this missing layer was part of its parent region. As the glomerular layer is the outermost layer of the main olfactory bulb, only voxels meeting this description were re-labeled.

#### S3.3 Manual annotation of newly reconstructed regions by experts

Manual expert annotation was performed using ITK-snap segmentation tool to complete the missing layers in the main olfactory bulb, specifically in the granular, inner plexiform, mitral, outer plexiform, and glomerular layers. Additionally, new labels were created for the granular and molecular layers across all 16 cerebellar lobules including the extended regions (see **Figure S8.1**). We based our delineation in the $\text{ARAva}_{\text{BBP}}$ $\text{Nissl}_{\text{COR}}$ on the distinct laminar structure of these regions. First, the missing annotated volume was completed with the outermost label from the ANNOTv3a, using a single view (coronal, sagittal, or horizontal) based on the presence of tissue in the Nissl volume. Second, the annotation of each layer was added in the same incidence as in the first step, based on the visual Nissl contrasts and/or textures. Third, the ANNOTv3a was corrected in each incidence to minimize discontinuities resulting in an accurate 3D annotated volume. Fourth, the ANNOTv3a was edited using 3D meshes (previews) for each layer to ensure there were no holes, resulting in a smooth volume. The goal of the third and fourth steps was to achieve a good compromise for each layer between maintaining consistency with the preexisting atlas, the aligned $\text{ARAva}_{\text{BBP}}$ $\text{Nissl}_{\text{COR}}$, and the new manual labels produced.

#### S3.4 Semi-automated method for producing the labels on the newly reconstructed regions

After having produced the $\text{AIBS}_{\text{BBP}}$ $\text{Nissl}_{\text{SAG}}$ and $\text{WAXH}_{\text{BBP}}$ $\text{Nissl}_{\text{HOR}}$ aligned to the $\text{ARAva}_{\text{BBP}}$ $\text{Nissl}_{\text{COR}}$ (see **Supplementary Material S7.2**), we averaged those three volumes to get better smooth contrasts between regions. This allowed us to design a semi-automated method composed of image processing techniques as well as minimal manual corrections (**Figure S3.4**) for identifying the labels in the extended part, as well as the granular and molecular layers in the cerebellum. A dedicated method was used for each region under consideration. Concerning the medulla, a simple process was set up as only a binary identification of the tissue was performed. This region underwent extensive manual processing, including smoothing and correction, to address the substantial noise present in the WAXH $\text{Nissl}_{\text{HOR}}$ medulla. These adjustments were necessary also to ensure consistency and continuity with ANNOTv3 from the AIBS. No additional expert annotation was produced for that region. For the main olfactory bulb, a progressive growing region process was designed, starting with the identification of the innermost region (granular layer, MOBgr) using Otsu thresholding, and ending with the outermost region (glomerular layer, MOBgl). The inner plexiform (MOBipl) as well as the mitral (MOBmi) layers were automatically estimated using a dilation with a 3 isotropic voxel kernel, in accordance with the ANNOTv3. Otsu thresholding made it possible to distinguish the outer plexiform layer (MOBopl) from the glomerular layer in the remaining unannotated tissue. After some smoothing operations, minimal manual corrections were applied for ensuring continuity between the new labels in the extended part and the ANNOTv3 for the main olfactory bulb layers. The method used for the cerebellum was different because it required identifying layers from scratch within each of its 16 lobules. We processed each of the lobules independently with the same method to provide a better precision in layer identification. Manual corrections were applied for ensuring continuity of the layers between lobules according to the ANNOTv3. Despite aligning and averaging 3 different brains, many artifacts of tearing, folding or missing tissue are still present in the data, especially in the cerebellum, which is why we had to perform manual artifact correction. Each of these corrections was performed using the Paxinos and Franklin paper atlas Fifth edition, as well as the expert delineations of the AIBS Nissl$_{SAG}$ on the web interface from the Allen Institute as a support.

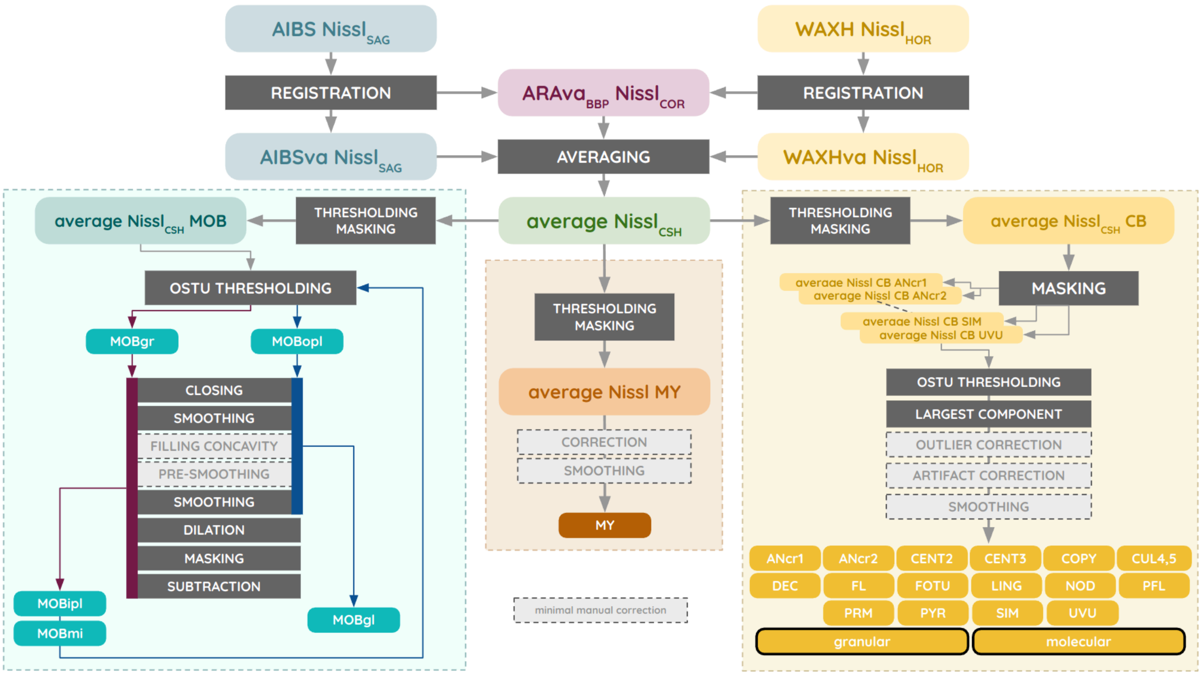


**Figure S3.4** | Semi-automated method for identifying the labels in the extended parts of the main olfactory bulb (MOB), cerebellum (CB), and medulla (MY), as well as the granular and molecular layers in each of the 16 lobules in the cerebellum. This method is applied on the average Nissl$_{CSH}$ derived from the three Nissl-stained volumes AIBS Nissl$_{SAG}$, WAXH Nissl$_{HOR}$ and ARAva$_{BBP}$ Nissl$_{COR}$.

#### S3.5 Assessment of the newly added labels with manual expert annotation

Visual assessment as well as Dice score calculation were performed on each of the extended/added layers in the main olfactory bulb and the cerebellum. The **Figure S3.5A** shows that the difference between the two versions, measured at the boundary between layers, rarely exceeds a thickness of 3 voxels. Most of the differences are between 0 and 1 voxel thickness. This tends to be confirmed by the Dice score for all regions in the **Figure S3.5B**, which is 0.97 on average for the olfactory bulb and the cerebellum, the lowest score being 0.94 obtained for the mitral layer of the main olfactory bulb. All the labels generated by the semi-automated method described in section S3.4 were validated by these high scores, indicating that our method accurately reproduced the expert labels on the anatomical data $ARAva_{BBP}$ $Nissl_{COR}$ with high precision.

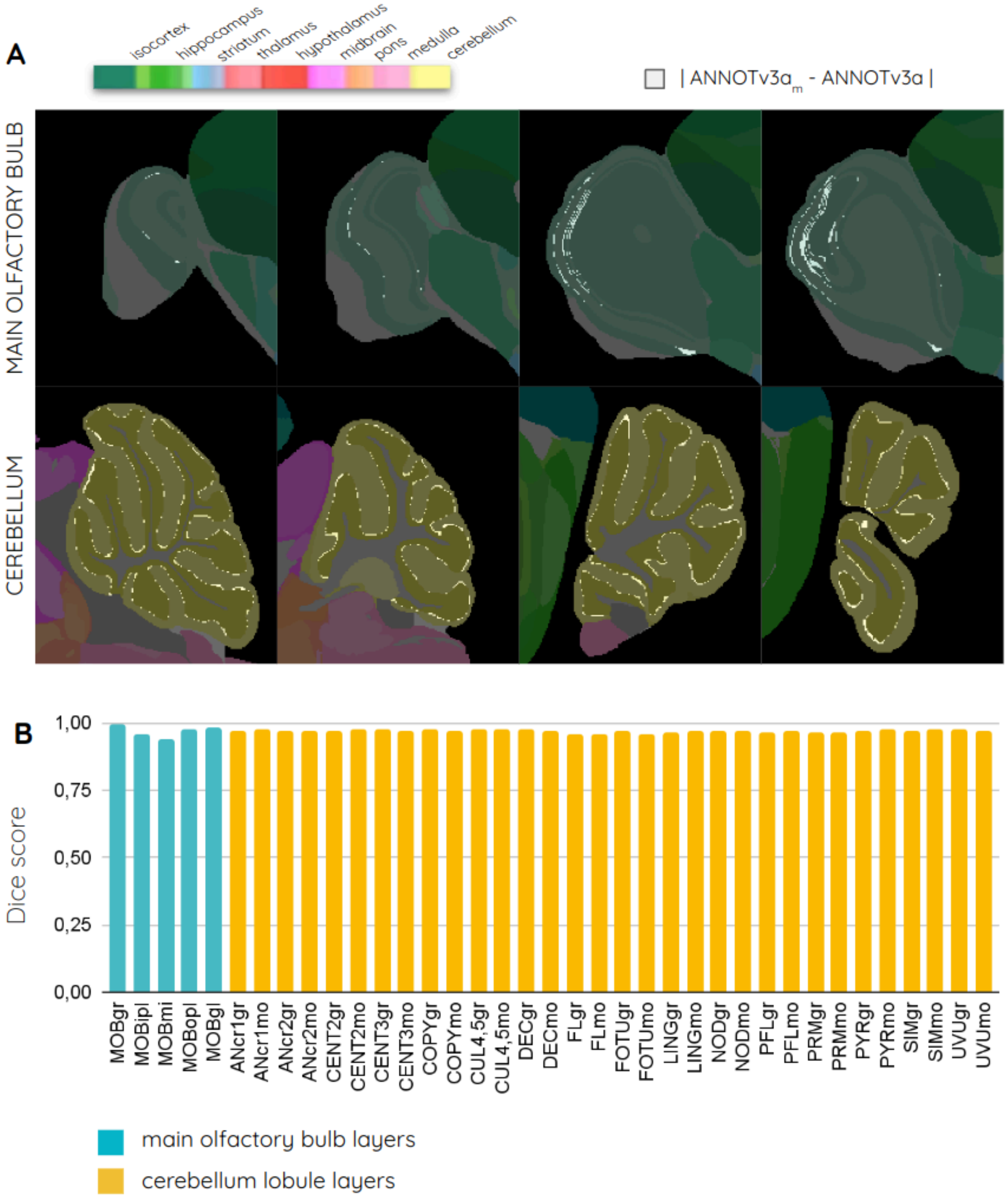


**Figure S3.5** | Comparison between the expert manual annotation ($ANNOTv3a_m$) and the annotation obtained by the semi-automated method presented in section S3.4 (ANNOTv3). (A) Absolute difference between the two compared labels, and (B) Dice scores evaluated for all the extended/added regions.

### S4 Hierarchy comparison between CCFv2 and CCFv3

The two latest versions of the annotation atlas provided by the Allen Institute CCFv2 (ANNOTv2) and CCFv3 (ANNOTv3) have different numbers of leaf regions (with non-zero voxel support), while they are based on the same hierarchy file. The ANNOTv3 shows a 14% reduction in the number of existing grey matter regions, while a 7% increase in the number of existing white matter regions, compared to ANNOTv2 (**Figure S4.1A-B**). Only 527 regions are therefore comparable when we consider the existing common regions in the two versions (**Figure S4.1C**).

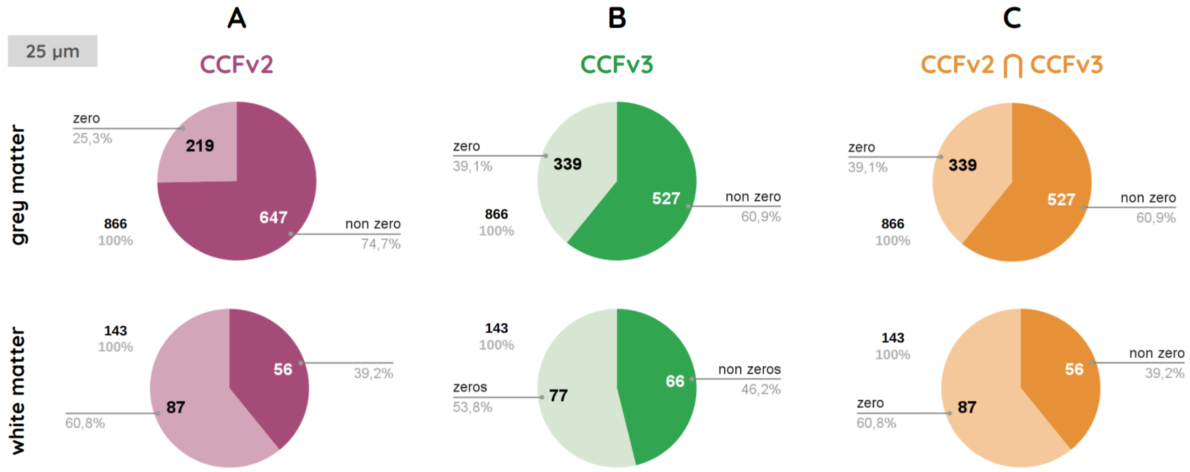


**Figure S4.1** | Existing regions (non-zero voxel support) in each annotation file. (A) CCFv2 and (B) CCFv3, as well as (C) their intersection, *i.e.* the common regions in the two versions.

Since not all regions have a direct counterpart between ANNOTv2 and ANNOTv3, we defined an intermediate level of ontology (referred to as 'parent'), outlined in **Figure S4.2**.

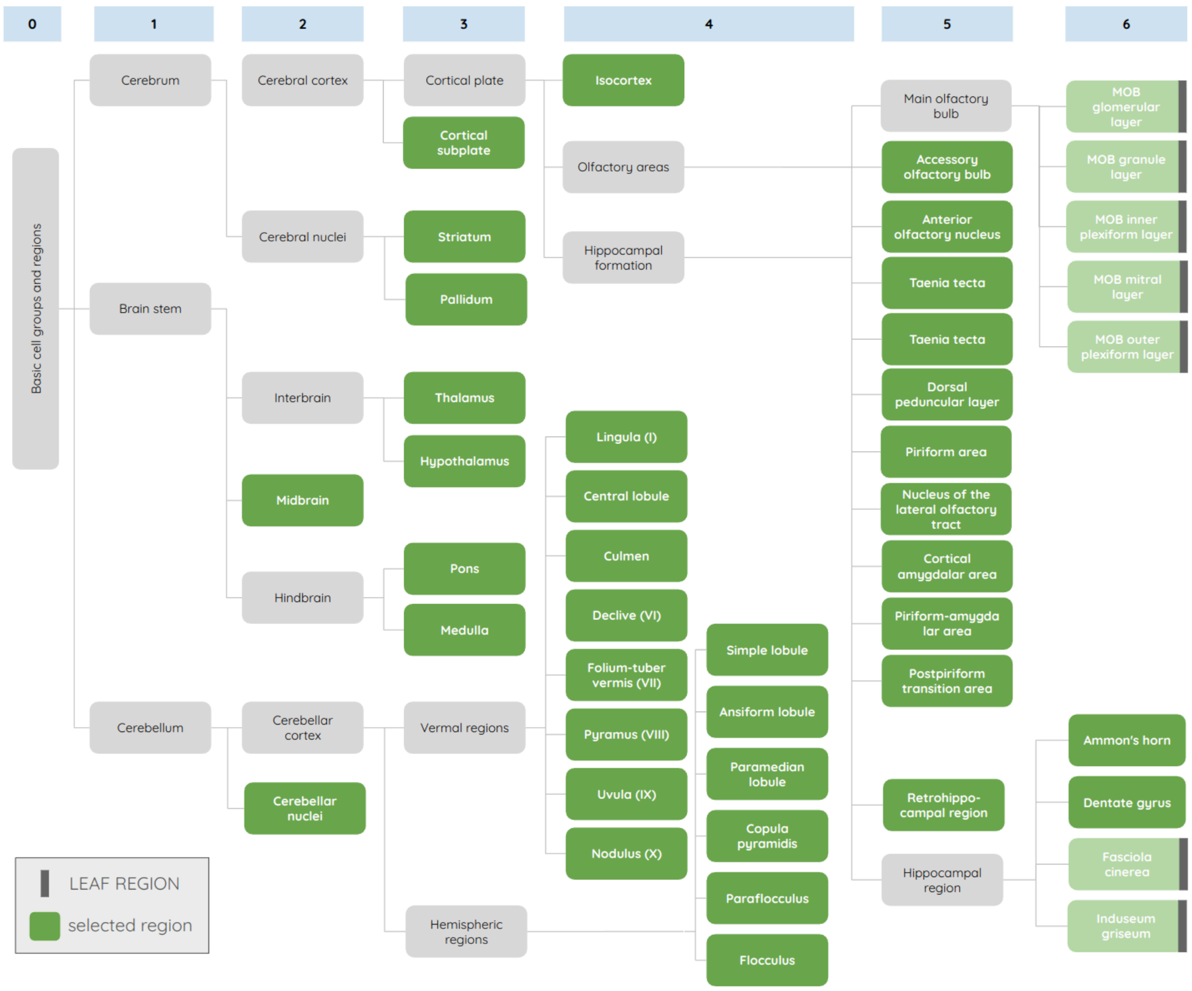


**Figure S4.2** | Overview of the second ontology level (parent) according to the different groups of regions in the hierarchy file for the region-based registration pipeline.

### S5 Slice-to-slice similarity

We provide detailed slices-to-slice Normalized Mutual Information (NMI) scores to evaluate the similarity between the template and the aligned $Nissl_{COR}$ in the three conventional incidences: coronal (**Figure S5.1A**), sagittal for the right hemisphere (**Figure S5.1B**), and horizontal (**Figure S5.1C**), comparing results before (ARA $Nissl_{COR}$) and after ($ARA_{BBP}$ $Nissl_{COR}$ after Step A5) alignment in the CCFv3.

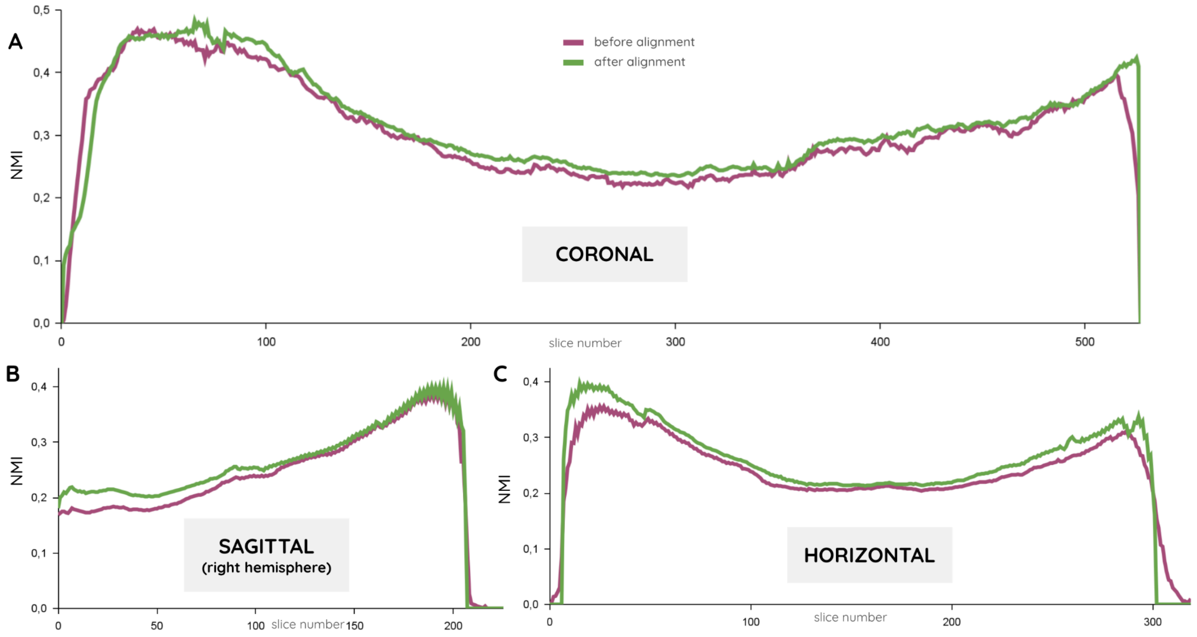


**Figure S5.1** | Slice-to-slice normalized mutual information estimated between each slice of the template and the ARA $Nissl_{COR}$ before (green) and $ARA_{BBP}$ $Nissl_{COR}$ after alignment (burgundy) in (A) the coronal incidence, (B) the sagittal incidence for the right hemisphere, and (C) the horizontal incidence.

### S6 Point identification in the ARA Nissl$_{COR}$

#### S6.1 Detailed method

We provide here the list of 29 fiducial points for assessing the quality of the alignment of the ARA Nissl$_{COR}$ in the CCFv3 (**Table S6.1** and **Figure S6.2**). All points were selected for their strategic placement at the boundaries between regions, clear 3D identifiability, and broad coverage of the main regions in the brain: olfactory areas, cerebellum, hippocampus, striatum, brainstem, thalamus, and cortex. We excluded points at the external border of the brain where it meets the background. Comprehensive documentation was provided for each point, enabling any operator to identify them without requiring specialized knowledge in neuroanatomy (**Table S6.1**). This documentation includes (1) the point's coordinates in the CCFv3 right hemisphere, (2) the region name and ID in the hierarchy it belongs to, (3) the preferred viewing plane for optimal identification, and (4) a detailed description of how the point was identified in 3D, with specific reference to its relative position along the rostro-caudal, dorso-ventral and medio-lateral axis. No leaf region in the hierarchy appears more than twice. Note that when multiple candidates appeared correct, the operator was asked to always select the one located at the center of the region's surface from that perspective.

Five different operators with various backgrounds and profiles (scientists or engineers, in biology, computational neuroscience, machine learning) identified a set of 29 fiducial points in each ARA Nissl$_{COR}$ volume before (ARA Nissl$_{COR}$) and after (ARA$_{BBP}$ Nissl$_{COR}$; after Step A5) alignment in the CCFv3. Operators were asked to give a score from 1 to 5 for each point to assess how straightforward it was to identify it in the Nissl tissue, defined as being: (1) very hard, (2) hard, (3) easy, (4) obvious, or (5) quite obvious. This confidence rate enabled us to remove the points for which this score was significantly low on average across all operators (< 2), meaning that the location of these specific points was not representative of their real anatomical location according to operators' evaluation. This way, we only kept points which were easy to recognize, eliminating any potential confusion in the identification process. Additionally, we removed points presenting inconsistent distances estimated to be greater than 1 millimeter, which indicates erroneous point identification, as no such high-intensity deformation was observed in the deformation field estimated at Step A5.

**Table S6.1** | Detailed description of the 29 fiducial points in the ANNOTv3.

| Acronym | Coordinates | Region name | ID | Incidence | Easy-to-understand description |
|---|---|---|---|---|---|
| OLF1 | [55, 164, 29] | Main olfactory bulb, inner plexiform layer | 228 | HORIZOTAL | The most lateral point of the medial side of the Main olfactory bulb, inner plexiform layer |
| OLF2 | [57, 138, 44] | Accessory olfactory bulb, granular layer | 196 | HORIZONTAL | The most rostral/proximal point of the Accessory olfactory bulb, granular layer |
| OLF3 | [94, 238, 33] | Main olfactory bulb, granule layer | 220 | SAGITTAL | The most ventral point of the Accessory olfactory bulb, granular layer |
| OLF4 | [52, 166, 13] | Main olfactory bulb, glomerular layer | 212 | HORIZONTAL | The most lateral point of the medial side of the Main olfactory bulb, glomerular layer |
| OLF5 | [22, 136, 15] | Main olfactory bulb, granule layer | 220 | HORIZONTAL | The most ventral point of the dorsal part of the Main olfactory bulb, granule layer |
| HIP1 | [301, 61, 61] | Field CA1, pyramidal layer | 407 | CORONAL | The most dorsal point of the Field CA1, pyramidal layer |
| HIP2 | [245, 97, 27] | Field CA3, pyramidal layer | 495 | HORIZONTAL | The most rostral point of the Field CA3, pyramidal layer |
| HIP3 | [333, 155, 119] | Field CA3, pyramidal layer | 495 | SAGITTAL | The center of the merging between the two parts of the Field CA3, pyramidal layer |
| HIP4 | [359, 127, 108] | Dentate gyrus, granule cell layer | 632 | SAGITTAL | The most caudal point of the dorsal Dentate gyrus, granule cell layer after the separation |
| HIP5 | [359, 156, 153] | Field CA1, pyramidal layer | 407 | SAGITTAL | The most caudal point of the Field CA1, pyramidal layer before the breaking of the circle it forms |
| STR1 | [253, 101, 94] | Caudoputamen | 672 | SAGITTAL | The most dorsal point of the Caudoputamen |
| STR2 | [204, 235, 99] | Caudoputamen | 672 | CORONAL | The most ventral point of the Caudoputamen |
| STR3 | [250, 230, 78] | Central amygdalar nucleus, medial part | 559 | CORONAL | The most medial point of the Central amygdalar nucleus, medial part |
| STR4 | [175, 264, 56] | Nucleus accumbens | 56 | HORIZONTAL | The most ventral point of the Nucleus accumbens |
| STR5 | [152, 216, 4] | Nucleus accumbens | 56 | SAGITTAL | The center point of the Nucleus accumbens between the two parts of the anterior commissure, olfactory limb just before the latter joins into one single peace |
| CB1 | [484, 195, 118] | Paramedian lobule, granular layer | 10681 | SAGITTAL | The most ventral point of the Paramedian lobule, granular layer |
| CB2 | [456, 49, 67] | Simple lobule, granular layer | 10672 | SAGITTAL | The most dorsal point of the Simple lobule, granular layer |
| CB3 | [501, 90, 93] | Crus 2, granular layer | 10678 | SAGITTAL | The most dorsal point of the Crus 2, granular layer |
| CB4 | [448, 259, 149] | Paraflocculus, granular layer | 10687 | SAGITTAL | The most ventral point of the Paraflocculus, granular layer |
| BS1 | [412, 195, 61] | Motor nucleus of trigeminal | 621 | SAGITTAL | The most dorsal point of the Motor nucleus of trigeminal |
| BS2 | [334, 165, 54] | Anterior pretectal nucleus | 215 | SAGITTAL | The most ventral point of the Anterior pretectal nucleus |
| BS3 | [429, 204, 13] | Abducens nucleus | 653 | SAGITTAL | The most dorsal point of the Abducens nucleus |
| BS4 | [435, 271, 74] | Facial motor nucleus | 661 | HORIZONTAL | The most lateral point of the Facial motor nucleus |
| BS5 | [395, 236, 13] | Tegmental reticular nucleus | 574 | SAGITTAL | The most caudal point of the Tegmental reticular nucleus |
| CTX1 | [239, 237, 117] | Basolateral amygdalar nucleus, anterior part | 303 | HORIZONTAL | The most rostral point of the Basolateral amygdalar nucleus, anterior part |
| CTX2 | [384, 88, 115] | Primary visual area, layer 6b | 305 | HORIZONTAL | The most caudal point of the Primary visual area, layer 6b |
| TH1 | [274, 111, 62] | Lateral dorsal nucleus of thalamus | 155 | SAGITTAL | The most dorsal point in the Lateral dorsal nucleus of thalamus |
| TH2 | [269, 131, 84] | Reticular nucleus of the thalamus | 262 | SAGITTAL | The most dorsal point in the Reticular nucleus of the thalamus |
| TH3 | [258, 153, 41] | Anteroventral nucleus of thalamus | 255 | SAGITTAL | The most caudal point of the Anteroventral nucleus of thalamus |

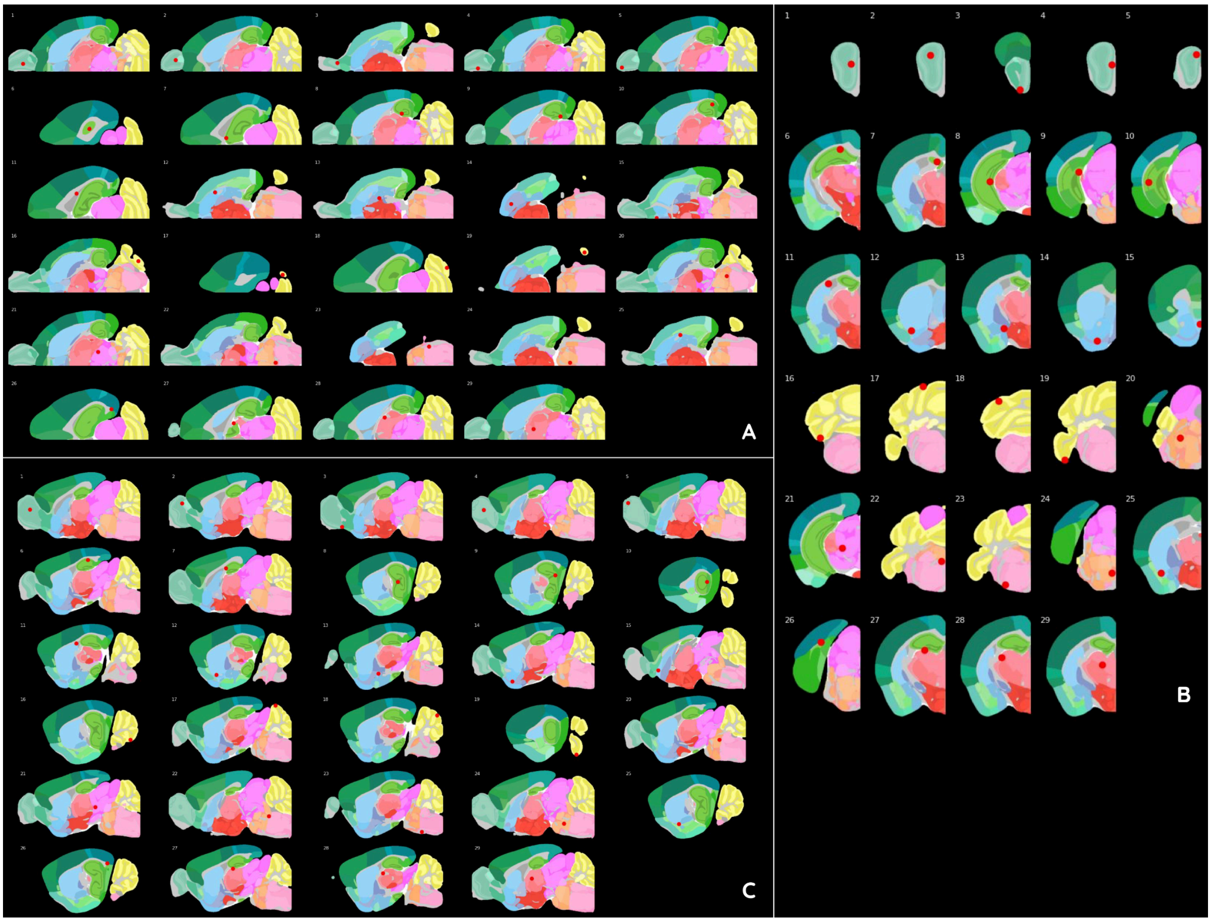


**Figure S6.2** | Plot of the 29 fiducial anatomical points of interest in the ANNOTv3 in the three conventional incidences (A) horizontal, (B) coronal, and (C) sagittal.

#### S6.2 Detailed Results

The detailed results of the Target Registration Error (TRE) calculated before and after the alignment are presented in **Figure S6.3**. Among the 29 fiducial points, three were considered to have insufficient confidence (STR2, STR3 and TH2; **Figure S6.3B**) and one was identified as an overlay (BS2 for operator 3: **Figure S6.3C**). Those three points (STR2, STR3, and BS2) were removed from the final average estimation, resulting in a 26 points dataset.

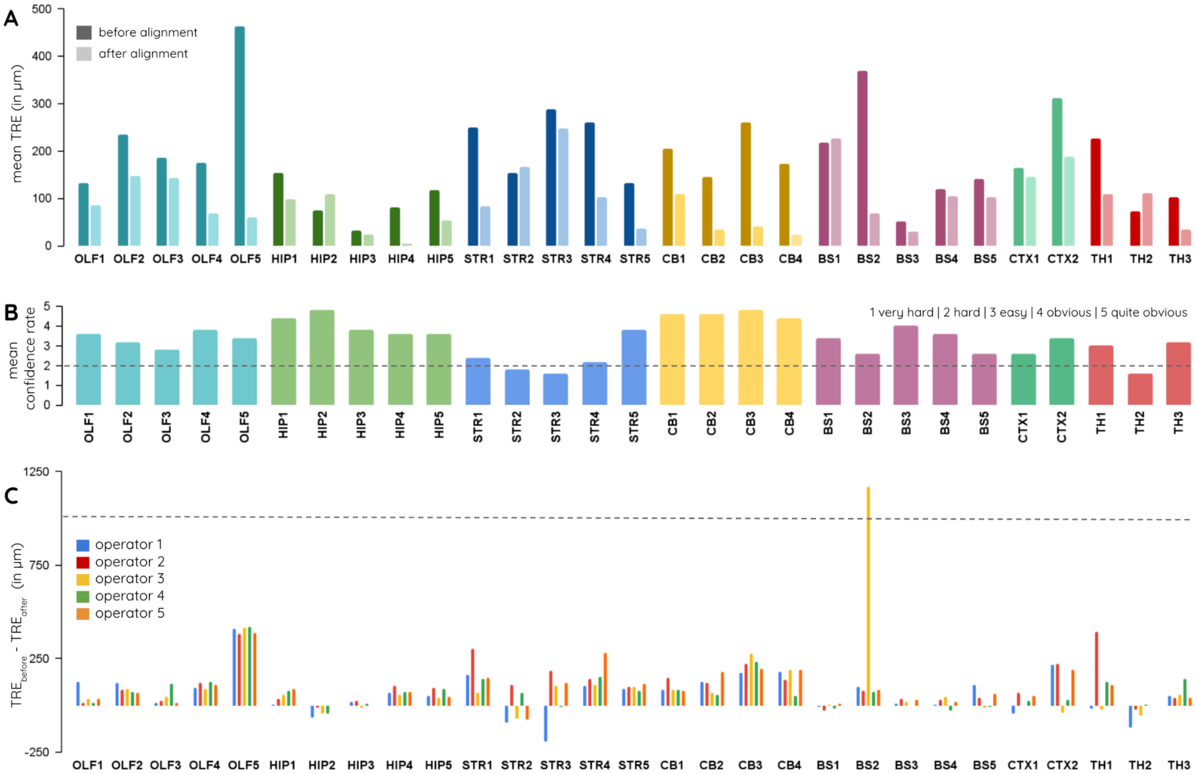


**Figure S6.3** | Detailed results for the point identification in the ARA Nissl$_{COR}$ before (ARA Nissl$_{COR}$) and after (ARA$_{BBP}$ Nissl$_{COR}$; after Step A5) alignment in the CCFv3. (A) Mean TRE calculated between the target (operator) and the reference (fiducial) point before (dark) and after (light) alignment, (B) mean confidence rate given by operators, and (C) TRE difference before and after the alignment for each of the 5 operators.

### S7 Detailed qualitative analysis of the alignment of the ARA Nissl$_{COR}$ in the CCFv3

#### S7.1 Aligning the ARA Nissl$_{COR}$ in the CCFv3

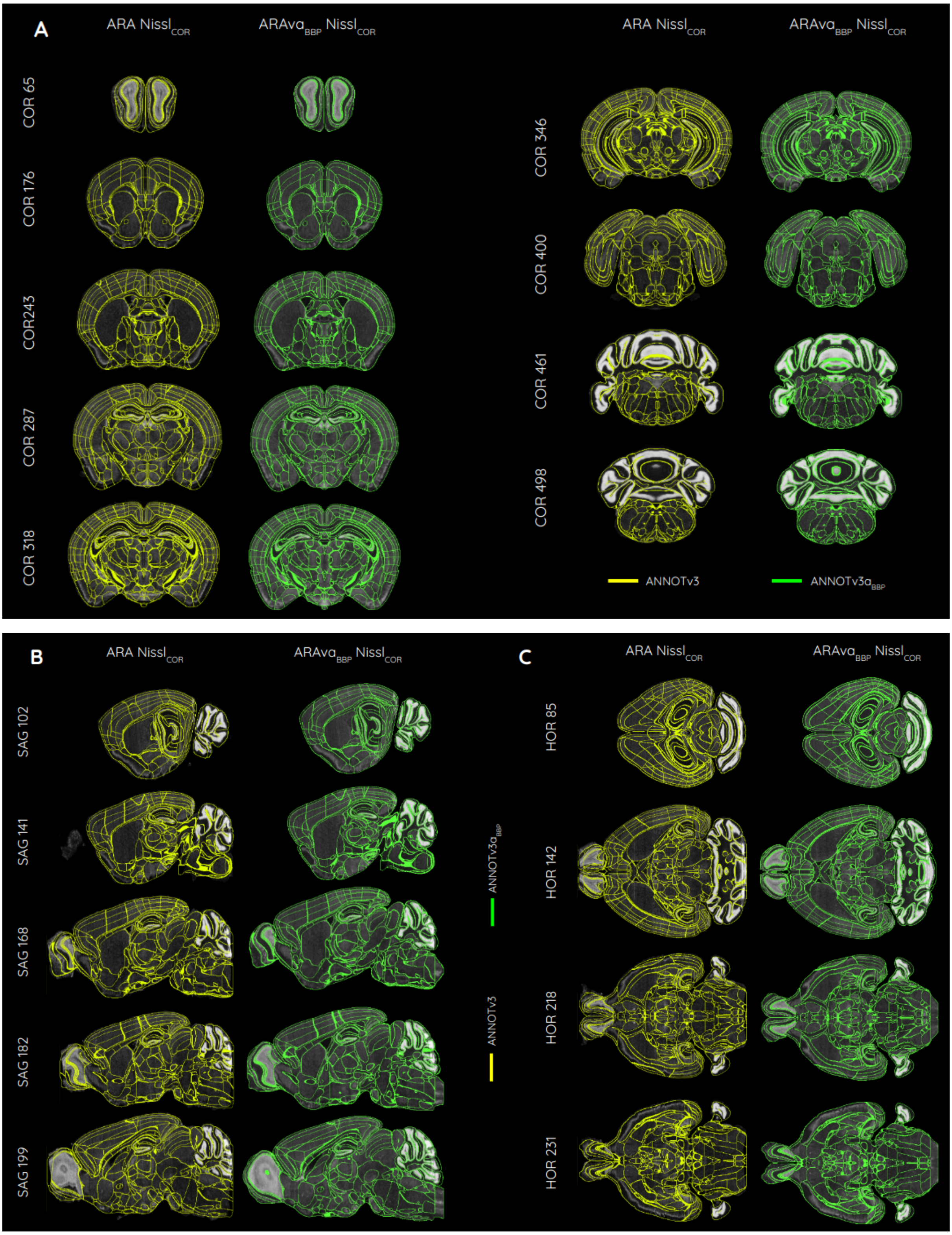


**Figure S7.1** | Overview of the superimposition of the annotations on the Nissl anatomy for the ARA Nissl$_{COR}$ (raw) with ANNOTv3 (yellow), and ARAva$_{BBP}$ Nissl$_{COR}$ (our alignment) with ANNOTv3a (green) in the (A) coronal, (B) sagittal, and (C) horizontal incidence.

#### S7.2 Aligning the AIBS Nissl$_{SAG}$ and WAXH Nissl$_{HOR}$ with the ARAva$_{BBP}$ Nissl$_{COR}$

The AIBS Nissl$_{SAG}$ and the WAXH Nissl$_{HOR}$ were both aligned to the ARAva$_{BBP}$ Nissl$_{COR}$ using the NiftyRefF3D registration algorithm, resulting into the AIBS$_{BBP}$ Nissl$_{SAG}$ and the WAXH$_{BBP}$ Nissl$_{HOR}$ (**Figure S7.2**).

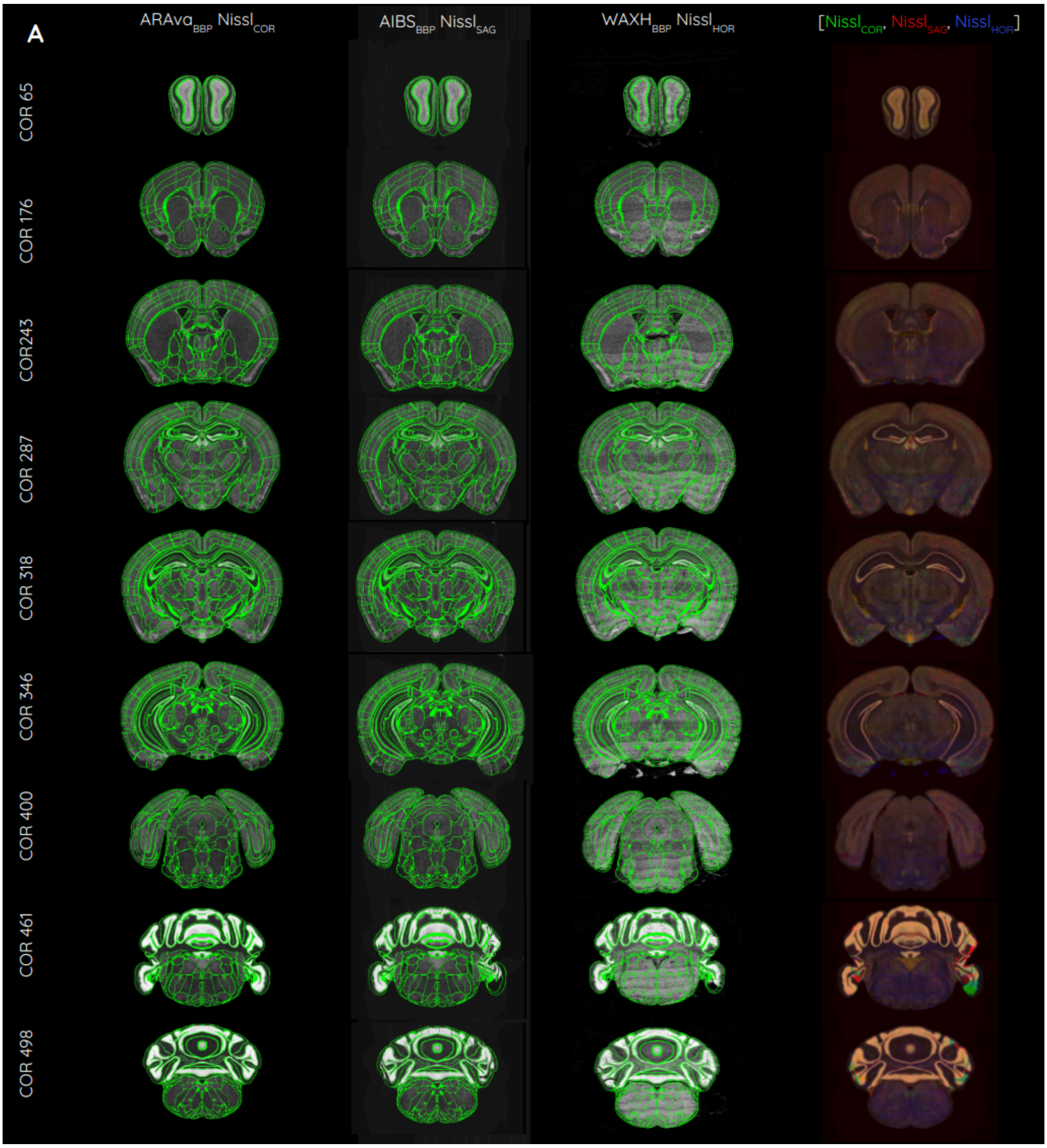


**Figure S7.2** | Overview of the three aligned volumes in the CCFv3 at 25 µm isotropic resolution: ARAva$_{BBP}$ Nissl$_{COR}$, AIBS$_{BBP}$ Nissl$_{SAG}$, and WAXH$_{BBP}$ Nissl$_{HOR}$ with the ANNOTv3a$_{BBP}$ boundaries overlayed (in green), plus, in the last column, the ARAva$_{BBP}$ Nissl$_{COR}$ is overlaid (in green), AIBS$_{BBP}$ Nissl$_{SAG}$ (in red), and WAXH$_{BBP}$ Nissl$_{HOR}$ (in blue) in the (A) coronal, (B) sagittal, and (C) horizontal incidence.

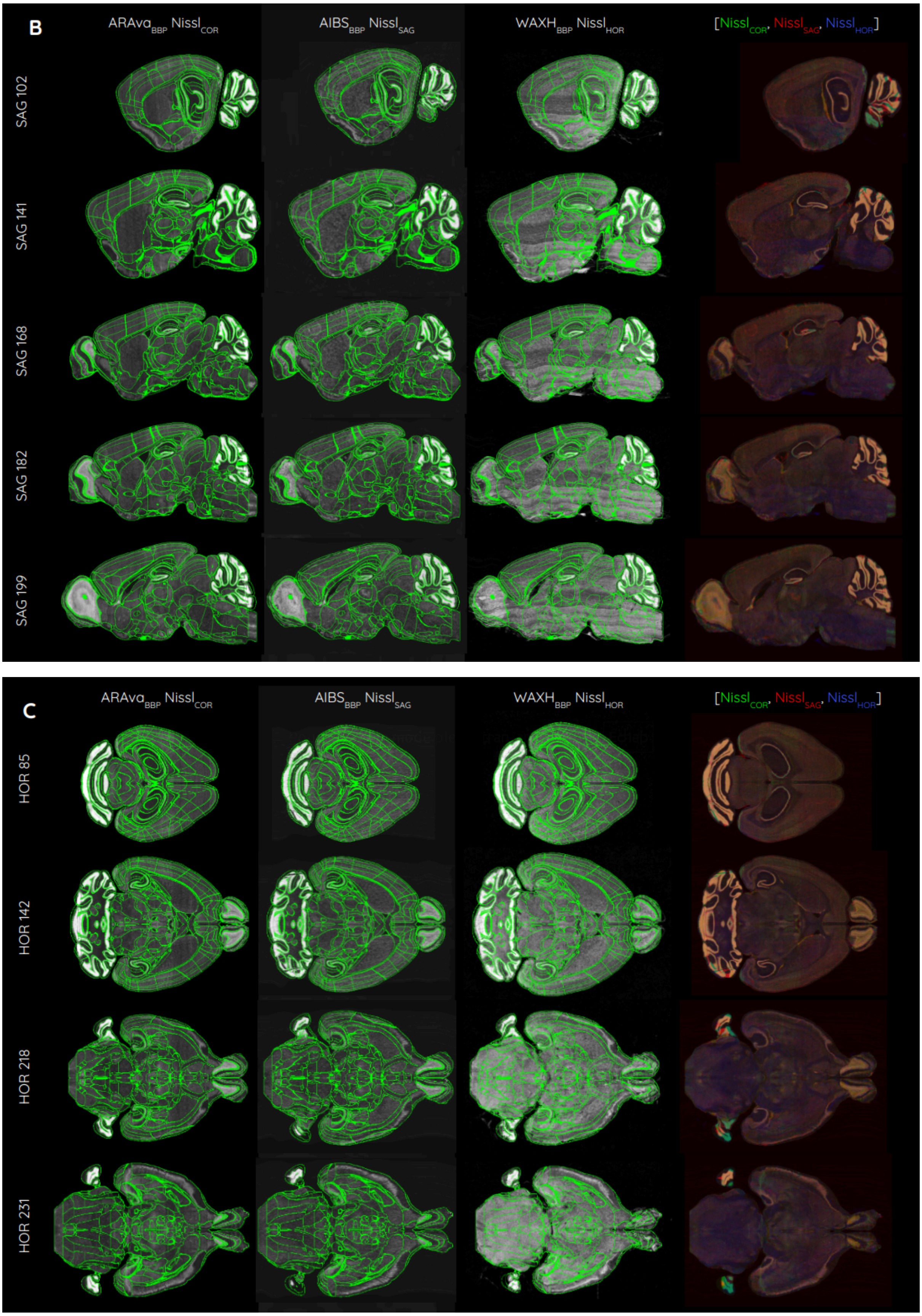
B
ARAva_BBP Nissl_COR
AIBS_BBP Nissl_SAG
WAXH_BBP Nissl_HOR
[Nissl_COR, Nissl_SAG, Nissl_HOR]
SAG 102
SAG 141
SAG 168
SAG 182
SAG 199
C
ARAva_BBP Nissl_COR
AIBS_BBP Nissl_SAG
WAXH_BBP Nissl_HOR
[Nissl_COR, Nissl_SAG, Nissl_HOR]
HOR 85
HOR 142
HOR 218
HOR 231

### S7.3 Producing an average Nissl template

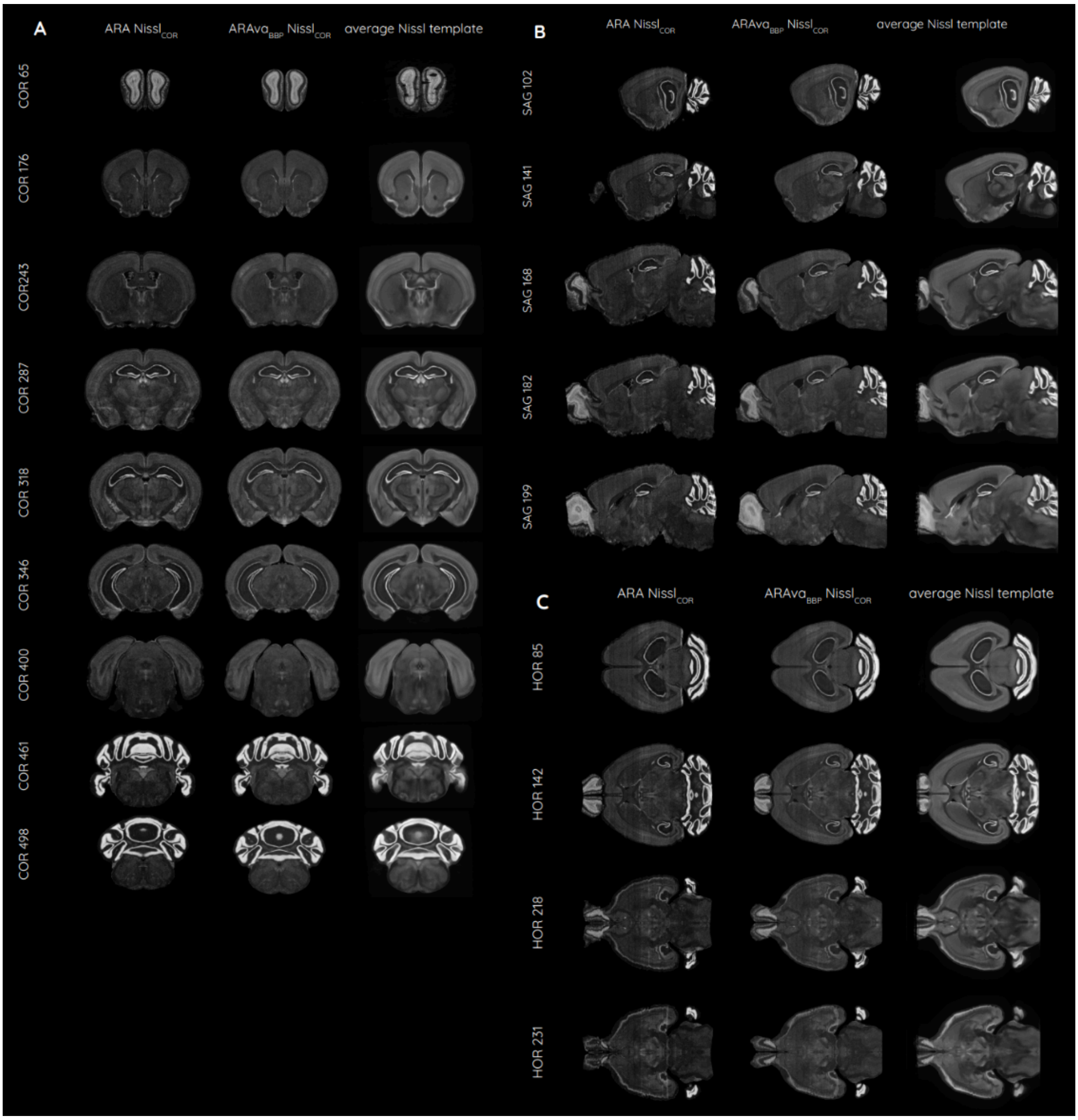


**Figure S7.3** | Overview of the three volumes ARA $Nissl_{COR}$ (raw), $ARAva_{BBP}$ $Nissl_{COR}$ (our alignment), and the average Nissl template (more than 86,000 registered coronal Nissl-stained slices from 734 mouse brains aligned on $ARAva_{BBP}$ $Nissl_{COR}$) in the (A) coronal, (B) sagittal, and (C) horizontal incidence.

### S8 Labels added in the cerebellum

We provide here an update on the cerebellum hierarchy in **Figure S8.1**, including 16 lobules: lingula (I), lobule (II), lobule (III), lobules (IV-V), declive (VI), folium-tuber vermis (VII), pyramus (VIII), uvula (IX), nodulus (X), simple lobule, crus 1, crus 2, paramedian lobule, copula pyramidis, paraflocculus, and flocculus. These 16 lobules each have 3 children defined in the hierarchy: granular layer, molecular layer and Purkinje layer. The ANNOTv3 provided by the Allen Institute does not include these regions. In the new version of the atlas we proposed, we produced the annotation for the two first ones (granular and molecular layers) at 25 µm isotropic resolution, plus the last one (Purkinje layer) at the junction between the granular and molecular layer of each lobule in the 10 µm isotropic resolution.

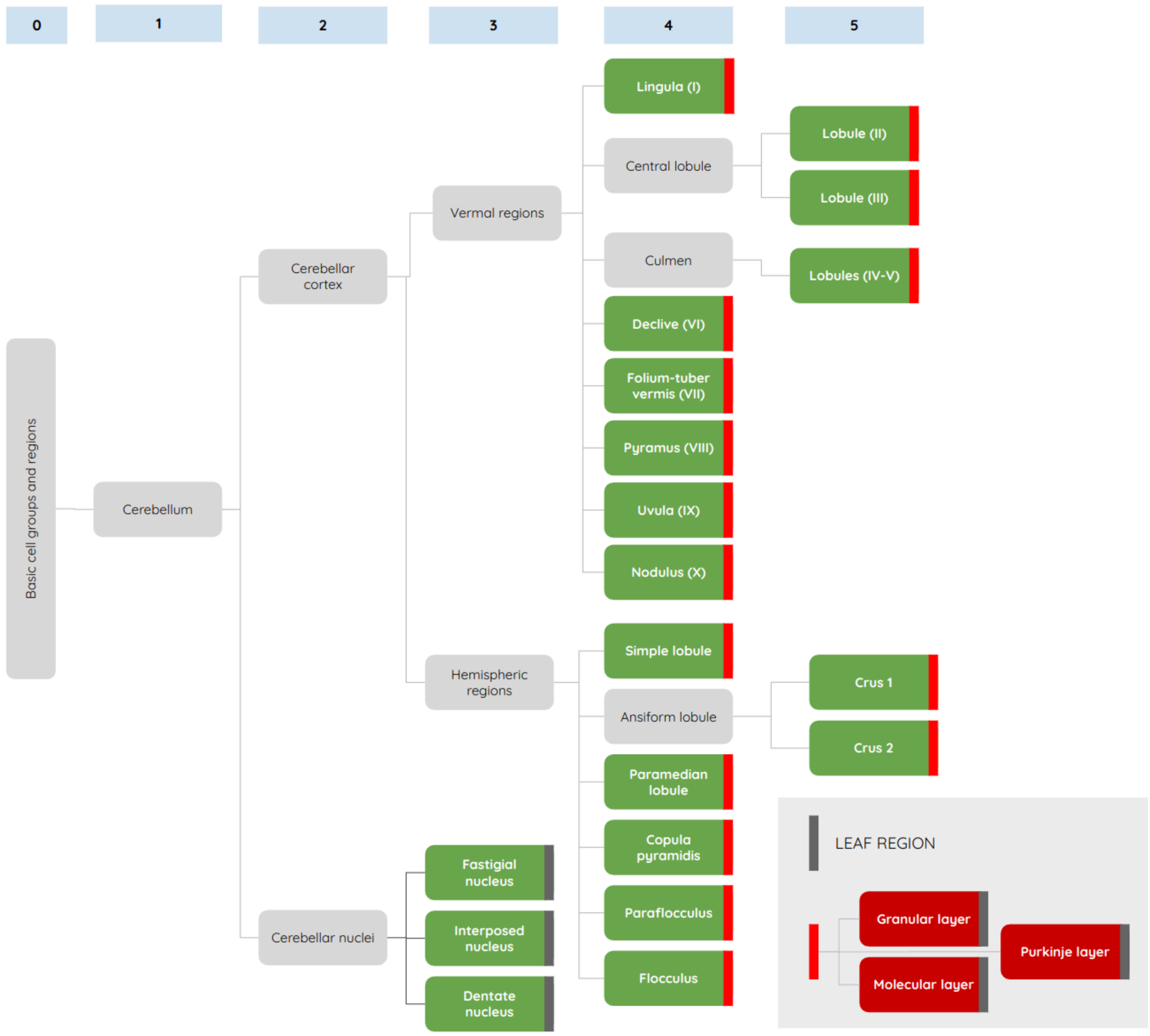


**Figure S8.1** | Cerebellum hierarchy, focusing on regions which are missing in the CCFv3 annotation (in red).

### S9 Number of voxels added the extended annotation

In this section, we detail the added and modified voxels resulting from the extended annotation (ANNOTv3a) process for specific regions such as the main olfactory bulb, the cerebellum, the medulla and the arbor vitae (**Table S9.1** and **Figure S9.2A**). Details are provided for layers of the main olfactory bulb and cerebellum in **Figure S9.2B-C**.

Added and modified voxels are defined as follows: (1) An added voxel is one that was unlabelled in the ANNOTv3 but received a label in the ANNOTv3a, (2) a modified voxel is one that was labeled in the ANNOTv3 but changed in the ANNOTV3a.

**Table S8.1** | Description of the number of voxels that were added and/or modified, as well as their proportion relative to the total volume of the main olfactory bulb (MOB), the cerebellum (CB), the medulla (MY) as well as the arbor vitae (at 25 µm isotropic resolution).

| | MOB | CB | MY | arbor vitae | TOTAL |
|---|---|---|---|---|---|
| **added** (nb. of voxels) | 30,870 | 161,026 | 244,784 | 302 | 436,982 |
| **added** (mm³) | 0.4823 | 2.5160 | 3.8248 | 0.0047 | 6.8278 |
| **added** (% of the region) | 2.6% | 4.5% | 11.0% | 0.1% | 1.3% |
| **modified** (nb. of voxels) | 345,440 | 3,291,636 | 592 | 0 | 3,637,668 |
| **modified** (mm³) | 5.3975 | 51.4318 | 0.0093 | 0.000 | 56.8386 |
| **modified** (% of the region) | 29.1% | 91.1% | 0.1% | 0.0% | 11.1% |
| **TOTAL** (nb. of voxels) | 376,310 | 3,452,662 | 245,376 | 302 | 4,074,650 |
| **TOTAL** (mm³) | 5.8798 | 53.9478 | 3.8340 | 0.0047 | 63.6664 |
| **TOTAL** (% of the region) | 31.7% | 95.6% | 11.1% | 0.1% | 12.4% |

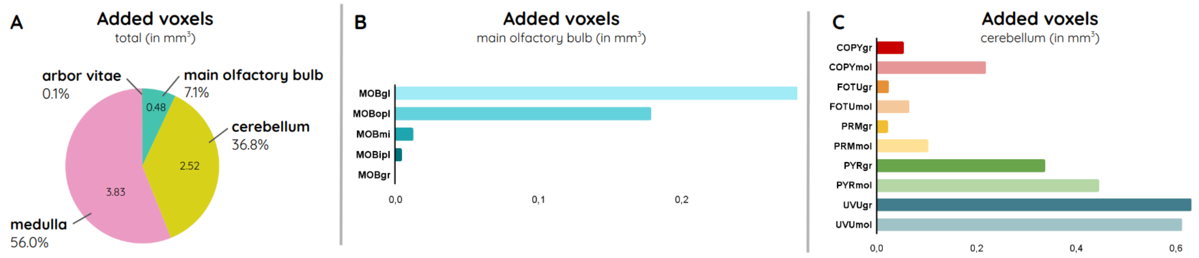


**Figure S9.2** | Detailed description of the added voxels (A) among all regions in the brain, (B) layers of the main olfactory bulb, and (C) cerebellar layers (at 25 µm isotropic resolution).

### S10 Aligning a gene expression dataset

The new extended ARAva$_{BBP}$Nissl$_{COR}$, aligned in the CCFv3, enabled the registration of gene expression dataset from the *in situ* hybridization (ISH) data portal at the Allen Institute (Lein et al., 2007; Ng et al., 2007). We selected a series of coronal slices and registered them to the extended and aligned ARAva$_{BBP}$ Nissl$_{COR}$ using the DeepAtlas tool (Krepl et al., 2021) and processed by the cell densities pipeline (Rodarie et al., 2022). Here are the genes we have selected: aldehyde dehydrogenase 1 family, member L1 (aldh1l1); 2',3'-cyclic nucleotide 3' phosphodiesterase (cnp); glial fibrillary acidic protein (gfap); myelin basic protein (mbp); S100 protein, beta polypeptide, neural (s100b); and the expression of glutamate decarboxylase 1 (gad67$_E$); parvalbumin (pv$_E$); somatostatin (sst$_E$); vasoactive intestinal polypeptide (vip$_E$). Registered coronal slices are presented in **Figure S10**.

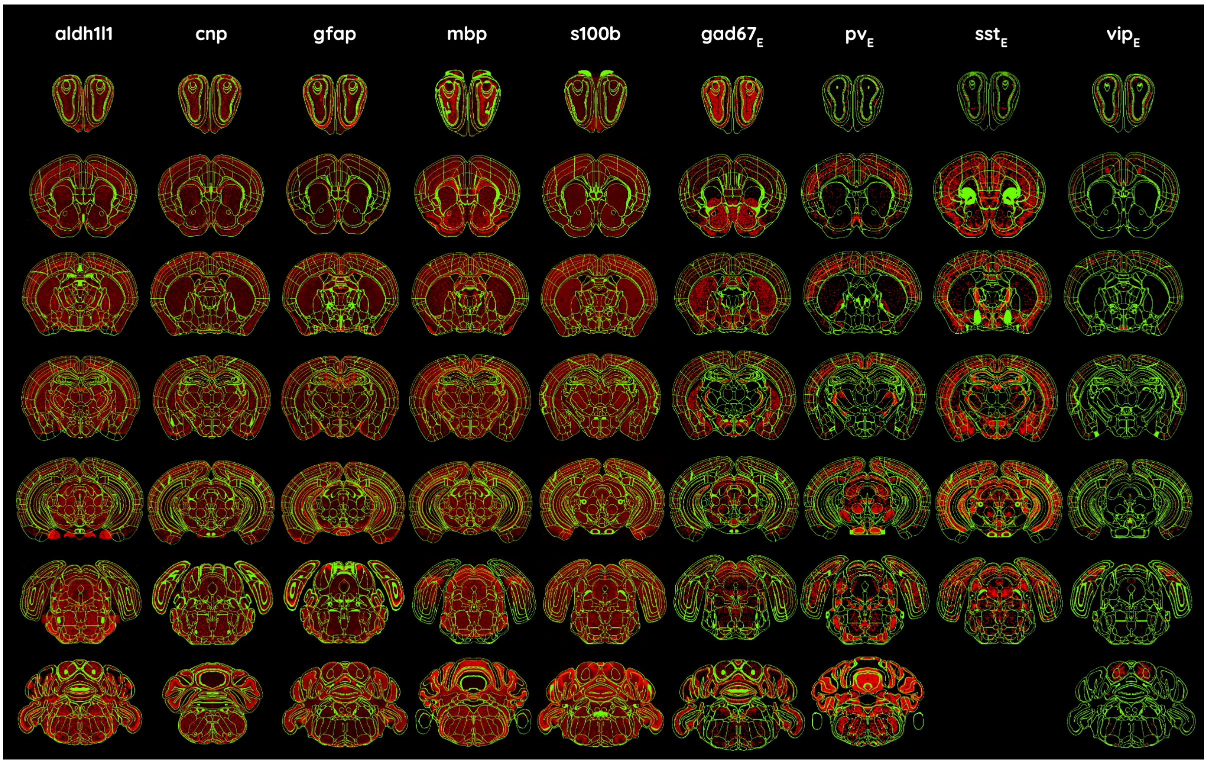


**Figure S10** | Coronal slices showing gene expression for nine different genes (in red) from the ISH data portal at the Allen Institute, with the CCFv3a$_{BBP}$ annotation overlaid (in green).

### S11 From *post mortem* to *in silico* cell modeling

The 3D Blue Brain atlas pipeline, utilizing our newly extended and aligned version, generated density maps for cells, neurons, inhibitory neurons, and excitatory neurons throughout the entire brain in the CCFv3. Several views of the produced data are shown in **Figure S11**.

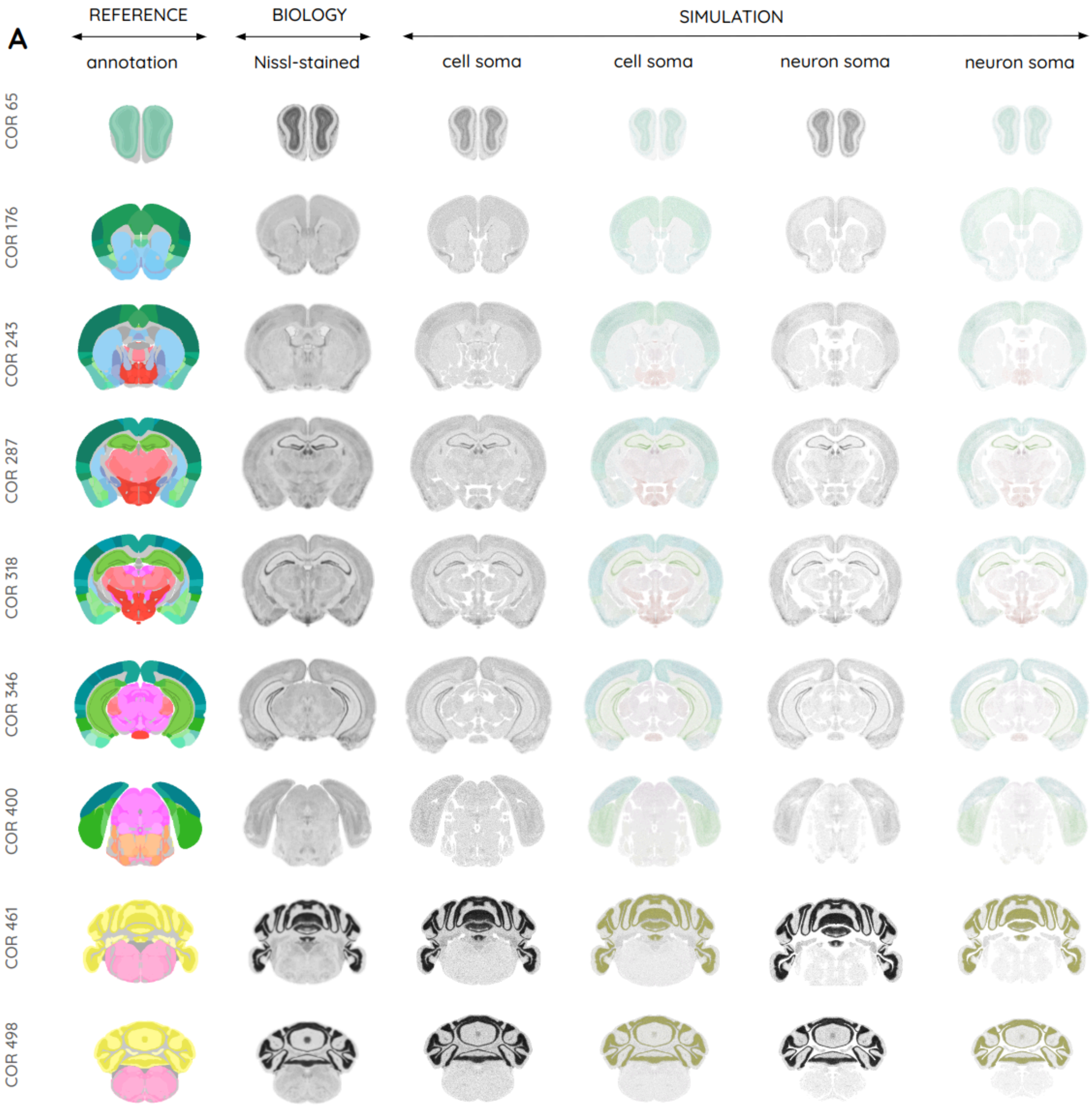


**Figure S11** | An overview of the simulated cell/neuron somas compared to real biological tissue ($ARAva_{BBP}$ $Nissl_{COR}$) from which it has been generated. These are merged with the $ANNOTv3a_{BBP}$ colors in the (A) coronal, (B) sagittal, and (C) horizontal views.

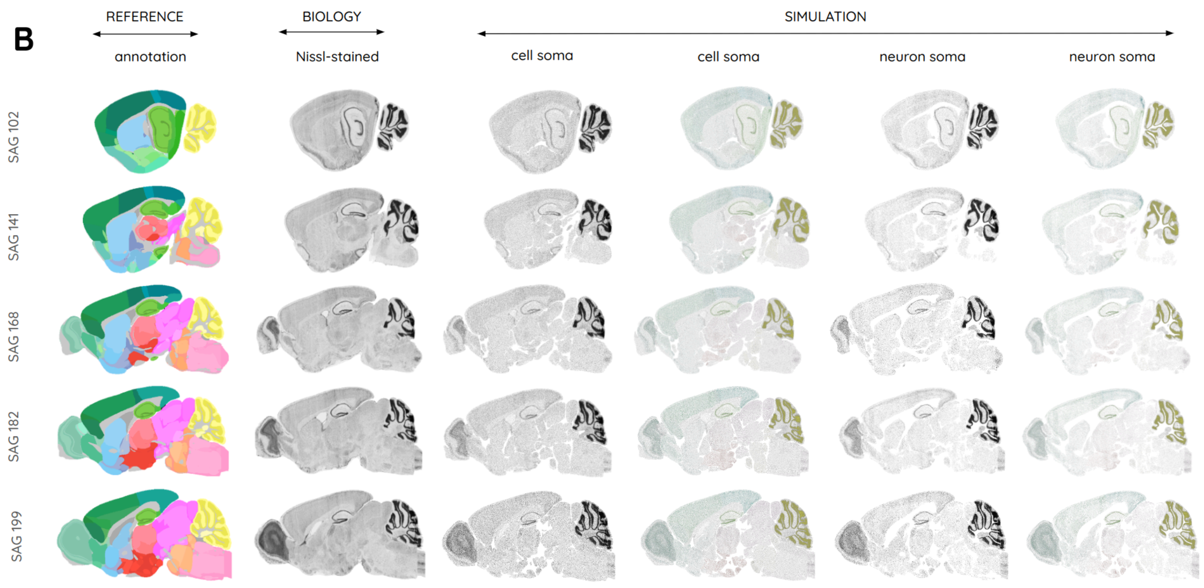
B
REFERENCE
annotation
BIOLOGY
Nissl-stained
SIMULATION
cell soma
cell soma
neuron soma
neuron soma
SAG 102
SAG 141
SAG 168
SAG 182
SAG 199

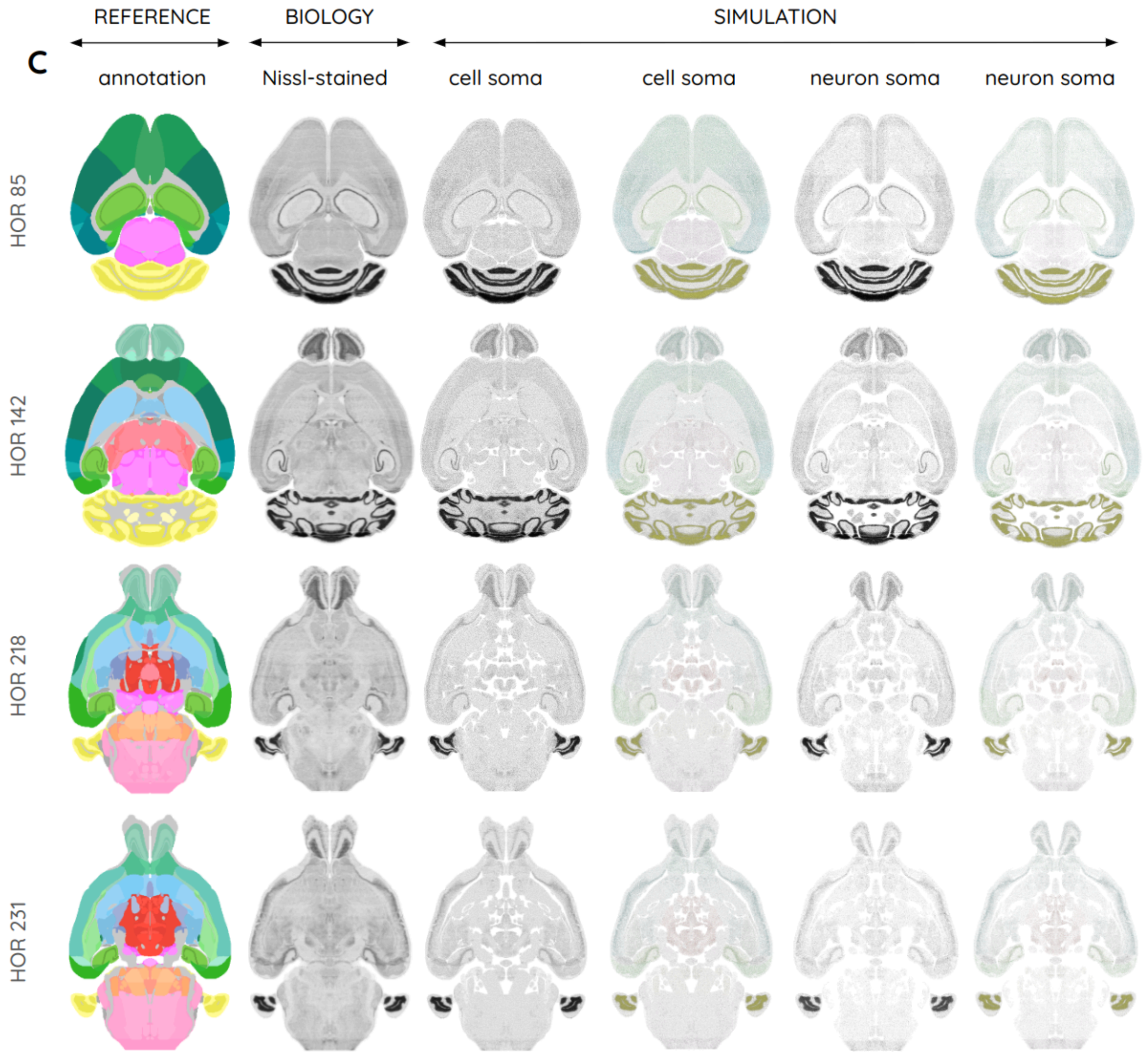
C
REFERENCE
annotation
BIOLOGY
Nissl-stained
SIMULATION
cell soma
cell soma
neuron soma
neuron soma
HOR 85
HOR 142
HOR 218
HOR 231
